# Phase-specific stimulation of prefrontal beta oscillations modulates distinct cognitive processes

**DOI:** 10.64898/2026.11.10.765583

**Authors:** Miles Wischnewski, Malte R. Güth, Ivan Alekseichuk, Alexander Opitz

## Abstract

Oscillatory phase has been proposed to organize neural information processing, but whether endogenous phase can be leveraged to causally shape human cognition remains unclear. Here, we tested whether repetitive transcranial magnetic stimulation (rTMS) delivered at specific phases of endogenous beta oscillations over the left dorsolateral prefrontal cortex modulates working memory and response inhibition. Participants received EEG phase-controlled, closed-loop rTMS delivered at either the peak or trough of their endogenous beta rhythm, using each participant’s dominant beta frequency (20-30 Hz). A conventional 20-Hz rTMS delivered independently of the beta phase served as an active control. Compared to this control and each other, peak-targeted rTMS improved response inhibition, whereas trough-targeted rTMS improved working memory. Both phase-specific behavioral effects were accompanied by corresponding increases in event-related potentials, which were not observed after conventional stimulation. Across participants, behavioral improvements correlated with stimulation induced electrophysiological changes. Finally, phase-specific rTMS induced distinct DLPFC network expansions that complemented the behavioral effects: peak stimulation enhanced posterior centro-parietal connectivity for inhibitory control, while trough stimulation expanded anterior prefrontal networks for working memory. These findings provide first evidence that opposite phases of endogenous prefrontal beta oscillations coordinate distinct cognitive and neural processes. This establishes prefrontal beta phase as a causal dimension for shaping executive function and as a target for prefrontal rTMS to modulate cognition and non-invasively probe phase dependent brain function.

## Introduction

The dorsolateral prefrontal cortex (DLPFC) is a key node in the brain’s cognitive control network, supporting essential executive functions like working memory and response inhibition (E. K. Miller & Cohen, 2001).The DLPFC is implicated in several cognitive operations and coordinates them by dynamically shifting activity across distinct, large-scale networks depending on the task (Cieslik et al., 2013; Menon & D’Esposito, 2022). Therefore, a key question is how activity within this region is organized to support different forms of task-relevant information. Working memory requires the maintenance of task-relevant information, whereas response inhibition requires the suppression of inappropriate responses. Beta oscillations (14-30 Hz) are particularly relevant in this context, as they have been linked to cognitive control, maintenance of task representations, and inhibition (Engel & Fries, 2010; Schaum et al., 2021; Swann et al., 2009). Accordingly, beta phase may provide a temporal structure that influences how the DLPFC and connected networks process cognitive information. Furthermore, beta might govern long-distance connections by interacting with slow-wave frequencies (<12 Hz) (Engel & Fries, 2010). However, it remains unknown whether phase-specific modulation of DLPFC beta activity can alter neural and behavioral responses during working memory or response inhibition. Therefore, the DLPFC provides a compelling target for testing whether oscillatory phase can shape prefrontal cognition.

Across the brain, cognition and behavior are encoded through the firing of individual neurons, as well as emergent oscillatory fields from synchronous firing of large neural populations, such as beta oscillations (Buzsáki, 2010; Fries, 2015). These signals are interdependently linked, with emergent fields modulating individual spike timing through mechanisms such as ephaptic coupling (Anastassiou et al., 2011; Fröhlich & McCormick, 2010). Neuronal firing is often biased toward specific phases of ongoing oscillations and has been implicated in various cognitive domains, including spatial orientation (O’Keefe & Recce, 1993), sequential memory (Siegel et al., 2009), working memory, and cognitive control (Daume et al., 2024). Together, these findings suggest that the coupling of oscillatory phase and spiking may provide a temporal structure for organizing information processing. However, most of this evidence comes from invasive recordings and is largely correlative, leaving open whether phase can be targeted non-invasively to causally shape cognition in healthy humans. Non-invasive, phase-locked transcranial magnetic stimulation (TMS) can address this limitation by probing causal relationships between brain oscillations and cognition (Thut et al., 2012).

Direct recordings of neural firing in the human prefrontal cortex are rare and largely restricted to surgical patients. As such, whether neural states reflected in oscillatory phase can drive cognitive function and behavior in the neurotypical brain remains unclear. Recent developments combining TMS with real-time electroencephalography (EEG) enable oscillatory phase to be targeted non-invasively. TMS can transiently perturb cortical activity, while EEG can track ongoing oscillations and estimate their phase in real time (Wischnewski et al., 2024). This allows stimulation to be delivered at predefined phases of an endogenous rhythm, providing a causal test of whether stimulation effects depend on oscillatory phase. Moreover, repetitive TMS (rTMS) locked to endogenous oscillatory phase can modulate subsequent brain responses (Zrenner et al., 2018), raising the possibility that phase-specific rTMS may also influence neural and behavioral responses measured during cognitive tasks. Recently, phase dependencies observed using TMS-EEG have been validated against invasive recordings in humans and nonhuman primates, demonstrating that these non-invasive dependencies reflect underlying neural dynamics (Haigh et al., 2025). Thus, phase-specific TMS-EEG provides a non-invasive approach for probing whether endogenous oscillatory phase shapes cortical responses and cognition.

Opposing phases correspond to alternating states of neural excitation and inhibition (Buzsáki et al., 2012). Invasive human recordings show that population activity in the motor cortex is entrained to the phase of the beta rhythm during rest, which diminishes during task execution (K. J. Miller et al., 2012). This suggests that beta phase defines precise temporal windows for gating motor output. In the prefrontal cortex, cognitive functions, such as response inhibition, critically depend on these windows of fluctuating excitability reflected in the phase of prefrontal beta oscillations (Schaum et al., 2021; Swann et al., 2009). Therefore, we hypothesize that rTMS delivered at distinct phases will differentially engage prefrontal networks, resulting in phase-specific network engagement and behavioral after-effects.

Conventional prefrontal high-frequency (20 Hz) rTMS has been used to modulate cognitive and clinical outcomes, but it is typically applied without considering the individual beta rhythm and the phase of ongoing oscillation. If beta phase is functionally relevant for working memory and response inhibition, behavioral changes induced by rTMS should depend on the phase at which stimulation is delivered. In the present study, we therefore applied closed-loop rTMS to the left DLPFC, targeting either the peak or trough of each participant’s endogenous beta rhythm recorded over the stimulation site. Each rTMS train’s inter-pulse interval was tailored to match the individual participants’ dominant resting beta frequency over the DLPFC. Conventional 20 Hz rTMS, delivered independent of the ongoing beta phase, served as a control condition. We hypothesized that beta phase specific rTMS would improve behavioral responses during working memory and response inhibition compared with conventional rTMS. To assess the task-related neural signatures, we analyzed event-related potentials (ERPs). By comparing peak and trough stimulation, we further tested whether different phases of the endogenous beta cycle preferentially modulate neural and behavioral responses during working memory or response inhibition. Furthermore, to explore changes in large-scale network activity mediated by low-frequency (<12 Hz) oscillations (Buzsáki & Draguhn, 2004), we explored changes in low-frequency connectivity.

## Results

We implemented a phase-targeting rTMS protocol to test whether the timing of prefrontal stimulation relative to endogenous beta phase modulates cognition (Fig. 1). Ongoing EEG activity over the left DLPFC was recorded from F3, spatially filtered with the surrounding electrodes, and analyzed online using the Bayesian Temporal Prediction algorithm. The algorithm was trained on a 3 min resting EEG recording before stimulation and then used to predict each participant’s endogenous beta phase in real time. The control condition followed a conventional 20 Hz rTMS protocol (referred to as standard rTMS below), consisting of 2 s stimulation trains with 40 pulses, separated by 28 s intervals. The phase specific conditions used the same train structure, but stimulation was delivered at each participant’s individual beta frequency and targeted either the peak or trough of the predicted beta rhythm. Response inhibition was assessed with a Go/No-Go task during the 28 s intervals between stimulation trains, whereas working memory was assessed with 1-back, 2-back, and 3-back tasks before and after the intervention. We investigated stimulation effects on accuracy and ERPs during response inhibition and working memory, and additionally assessed functional connectivity at the sensor and source level before stimulation, between stimulation trains, and after stimulation.

**Figure 1.**
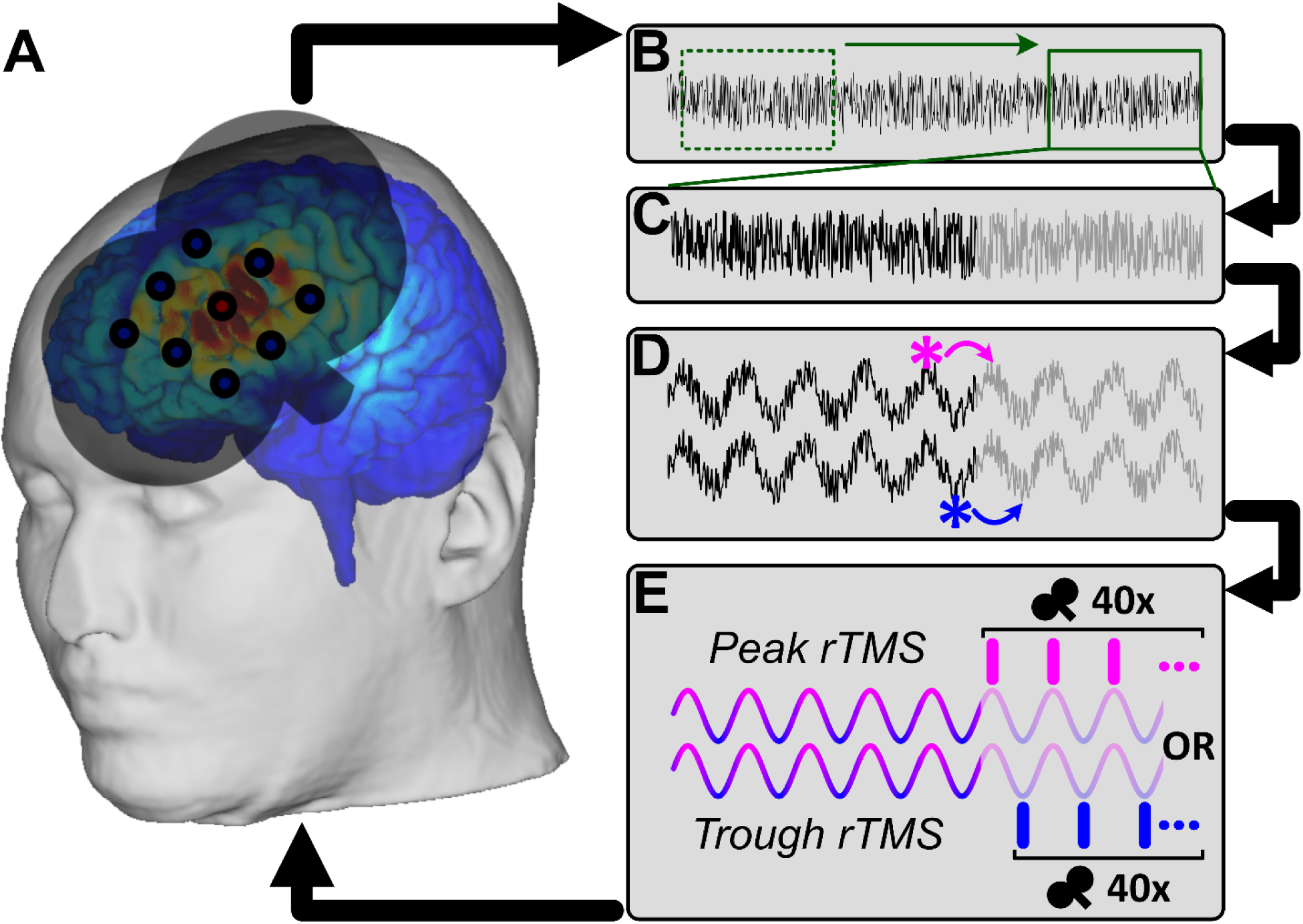
Phase-dependent beta-rTMS targeting of the left dorsolateral prefrontal cortex (DLPFC). A) EEG was recorded from 64 channels while rTMS was delivered over the left DLPFC (MNI coordinates [-40, 34, 28]), with online phase extraction centered on electrode F3 using Laplacian spatial filtering. (B–D) EEG data were streamed to MATLAB in real time, where the BTP algorithm identified and predicted the ongoing phase of prefrontal beta oscillations from individualized resting-state EEG parameters. Based on these predictions, stimulation bursts were timed to either the peak or trough of the beta rhythm. (E) In the phase-specific conditions, bursts of 40 pulses were delivered according to each participant’s individualized beta inter-cycle interval, whereas in the standard 20 Hz control condition pulses were delivered independent of the oscillatory phase.

### Modulation of inhibitory control

To assess the effects of rTMS on response inhibition, we quantified commission errors in the Go/No-Go paradigm (pressing the response button in NoGo trials) during NoGo trials, with fewer commission errors indicating better inhibitory control. Commission errors differed significantly between stimulation groups (F(2,63) = 9.84, p < 0.0001; Fig. 2A). Bonferroni-corrected post-hoc comparisons showed that significantly fewer errors were made in the peak rTMS group compared to the trough rTMS group (t(42) = 3.66, p_Bonf_ = 0.0021, d = 1.10) and standard rTMS group (t(42) = 4.00, p_Bonf_ = 0.0008, d = 1.21). No significant difference was observed between the trough and standard group (t(42) = 0.34, p_Bonf_ ≈ 1, d = 0.10). Also, no significant difference in reaction time was observed between groups (F(2,63) = 0.50, p = 0.6066), suggesting that better performance with peak rTMS cannot be explained by a difference in the speed-accuracy trade-off between experimental groups.

**Figure 2.**
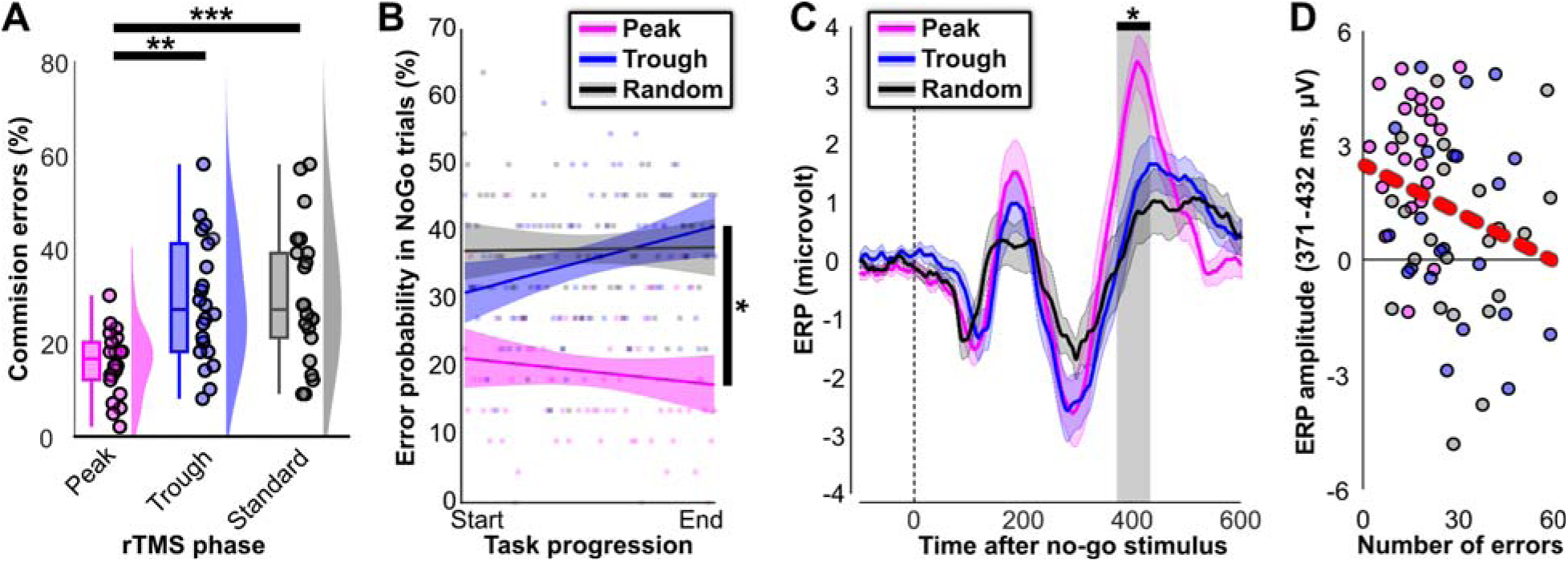
Modulation of inhibitory control with phase-specific beta rTMS. A) During peak-beta rTMS significantly fewer commission errors occurred compared to trough-beta and phase-unspecific 20 Hz rTMS. B) A scatter plot of the commission error probability from throughout out the task shows a significant interaction between peak-an trough-beta rTMS. While participants are less likely to make errors at the end of the task, compared to the beginnin in the peak condition, the opposite is observed for the trough condition. C) Event-related potentials after no-go stimuli for the three conditions. Noticeably, cluster-based permutation testing revealed a significant difference between 372-435 ms, corresponding to the P3 component. During peak beta-rTMS the P3 component was significantly larger compared to the trough-stimulation and control group. D) P3 amplitude per participant was significantly correlated t performance. Specifically, larger P3 amplitudes were associated with fewer commission errors. In all subplots significance is indicated as *p<0.05, **p<0.01, ***p<0.001.

To further explore how rTMS dynamically modulated behavior over time, we performed a multiple linear regression on the averaged commission errors over time (Fig. 2B). An interaction between trial number and the stimulation groups was found (t(234) = 2.54, p = 0.011), showing that errors gradually reduced in the peak rTMS condition, while errors gradually increased in the trough rTMS condition. Together the results imply that peak rTMS improved inhibitory control in comparison to the other two interventions.

### Modulation of electrophysiological components in Inhibitory control

To test whether the behavioral improvement in inhibitory control was accompanied by changes in task related brain activity, we compared NoGo event related potentials (ERPs) between stimulation groups. ERPs were analyzed separately over frontal and posterior midline regions using cluster based permutation testing across the post stimulus time window. Cluster-based permutation testing showed a significant difference in NoGo ERP signals in the frontal midline ROI between conditions between 371 and 432 ms after stimulus presentation (F_Clus_ = 1965.75, p = 0.0378; Fig 2C). ERP of each ROI channel separately is shown in Supplementary Fig 1,2. This observation is in line with previous research that suggested that NoGo modulation occurs in frontocentral regions, reflected by a P3 component (Albert et al., 2013). Bonferroni-corrected post hoc comparisons of the selected time window showed a significant difference between the peak and trough (t(42) = 2.97, p_Bonf_ = 0.0148, d = 0.90) and the peak and standard condition (t(42) = 3.71, p_Bonf_ = 0.0018, d = 1.12), but not between the trough and standard condition (t(42) = 0.56, p_Bonf_ ≈ 1, d = 0.17). Furthermore, we found a significant negative association between the number of errors and the P3 amplitude (r = -0.24, p = 0.0482), suggesting that participants who performed better also had larger component amplitudes (Fig. 2D). No significant ERP modulation was observed in the posterior midline ROI (p = 0.1938, Supplementary Fig 3). Together, these findings indicate that peak rTMS improved inhibitory control in comparison to trough and standard rTMS, and that this behavioral effect was accompanied by stronger inhibition related frontal activity. The association between both measures suggests that phase specific stimulation did not merely change task performance, but modulated neural processes relevant for inhibitory control.

### Modulation of working memory

To assess working memory, participants completed 1-back, 2-back, and 3-back tasks before and after the stimulation intervention. We quantified stimulation-induced changes in performance as the pre-to-post change in d-prime, with higher values indicating improved working memory accuracy. Pre-to-post changes in d-prime differed significantly between stimulation groups (F(2, 63) = 9.25, p = .0003, ηp² = .23). Despite the absence of a significant interaction effect between the TMS condition and the levels of the N-back (F(4, 126) = 1.49, p = .2102, ηp² = .05) follow-up one-way ANOVAs suggested the effect to be more pronounced for the 2-back (F(2, 63) = 8.00, p = .0008, ηp² = .21) and 3-back (F(2, 63) = 8.93, p = .0004, ηp² = .22), compared to the 1-back (F(2, 63) = 1.98, p = .1463, ηp² = .06). Bonferroni-corrected pairwise comparisons showed that, for the 2-back, d-prime increased more in the trough rTMS group compared to the peak rTMS group (t(42) = 2.97, p_Bonf_ = .0150, d = 0.90) and the standard rTMS group (t(42) = 3.77, p_Bonf_ = .0015, d = 1.14). Similarly, for the 3-back, d-prime increased more in the trough rTMS group than in the peak rTMS group (t(42) = 2.75, p_Bonf_ = .0264, d = 0.84) and the standard rTMS group (t(42) = 3.87, p_Bonf_ = .0011, d = 1.17).

For reaction time, there was no significant main effect of stimulation condition on pre-to-post change (F(2, 63) = 0.50, p = .6060, ηp² = .02), and no interaction between TMS group and working memory load (F(4, 124) = 1.38, p = .2450, ηp² = .04). Follow-up one-way ANOVAs likewise showed no condition effects for the 1-back (F(2, 63) = 0.41, p = .6654, ηp² = .01), 2-back (F(2, 63) = 1.84, p = .1678, ηp² = .06), or 3-back (F(2, 63) = 0.06, p = .9416, ηp² < .01). Together, these results indicate that trough rTMS selectively improved working memory accuracy, particularly under higher memory load, without evidence for a speed-accuracy trade-off.

### Modulation of electrophysiological components in Inhibitory control

To test whether the working memory improvement was accompanied by changes in task-related brain activity, we analyzed pre-to-post changes in N-back event-related potentials (ERPs). ERPs were analyzed separately for the 1-back, 2-back, and 3-back tasks over posterior and frontal midline regions using cluster-based permutation testing. Cluster-based permutation testing showed significant condition-dependent differences in pre-to-post rTMS changes in N-back ERP signals in the posterior midline ROI. For the 2-back task, a significant positive cluster was observed between 234 and 584 ms after stimulus presentation (F_Clus_ = 4824.34; p = 0.0002; Fig. 3E). Bonferroni-corrected post hoc comparisons of the selected time window showed a significant difference between the peak and trough condition (t(42) = 3.90, p_Bonf_ = 0.0010, d = 1.17) and between the trough and standard condition (t(42) = 2.73, p_Bonf_ = 0.0273, d = 0.82), but not between the peak and standard condition (t(42) = 1.65, p_Bonf_ = 0.3167, d = 0.50). Similarly, for the 3-back task, a significant positive cluster was observed between 246 and 434 ms (F_Clus_ = 2067.47; p = 0.0082; Fig. 3F). Post hoc comparisons showed a significant difference between the trough and standard condition (t(42) = 3.27, p_Bonf_ = 0.0064, d = 0.99), but not between peak and trough (t(42) = 2.04, p_Bonf_ = 0.1441, d = 0.61) or peak and standard (t(42) = 1.46, p_Bonf_ = 0.4541, d = 0.44). No significant posterior ERP modulation was observed for the 1-back task (p = 0.7776; Fig. 3D). The effects across all channels are shown in Supplementary Fig. 4-10. No significant clusters were observed in the frontal ROI (all p ≥ 0.2561; Supplementary Fig. 4), consistent with previous work suggesting that N-back-related ERP modulation is most pronounced over central-parietal regions (Scharinger et al., 2017).

**Figure 3.**
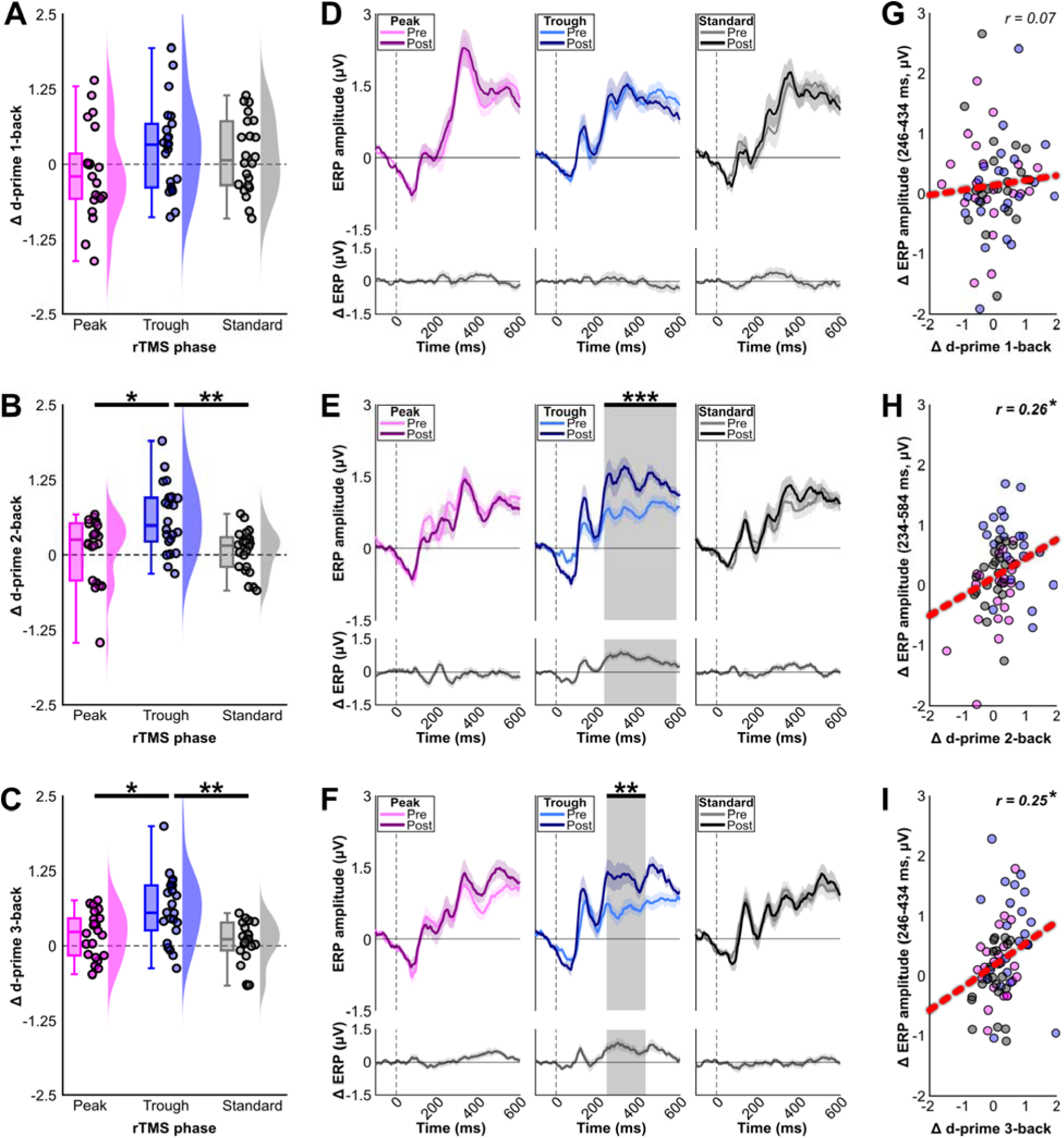
Modulation of working memory with phase-specific beta rTMS. Posttest - pretest d-prime scores were calculated for the A) 1-back, B) 2-back, and C) 3-back task. Trough-beta rTMS significantly increased d-prime more than peak-beta and phase-unspecific 20 Hz rTMS in the 2-back and 3-back, but not in the 1-back task. D) Event-related potentials after stimulus presentation were computed before and after rTMS for the D) 1-back, E) 2-back, an F) 3-back task. Cluster-based permutation testing revealed a significant difference in ERP amplitude from pre-t post-stimulation with trough-beta rTMS, but not with peak-beta and phase-unspecific rTMS. Significant clusters wer observed between 234 and 584 ms for the 2-back task, and between 246 and 434 ms for the 3-back task, with no differences for the 1-back task. The ERP post-pre difference clusters were correlated with post-pre behavioral changes for the G) 1-back, H) 2-back, and I) 3-back task. Because no significant cluster was observed for the 1-back task, ERP amplitudes for the 1-back correlation were extracted from the temporal overlap of the significant 2-back and 3-back windows, 246–434 ms. For both the 2-back and 3-back a significant correlation was observed, wit increasing differences in ERP amplitudes from pre to post rTMS being associated with increasing performance from pre to post rTMS. In all subplots significance is indicated as *p<0.05, **p<0.01, ***p<0.001.

We next tested whether stimulation-induced ERP changes were related to changes in working memory performance. Significant positive associations were found between posterior ERP amplitude changes and changes in d-prime. For the 2-back task, larger ERP changes were associated with larger improvements in d-prime (r = 0.264, p = 0.0319; Fig. 3H). A similar association was observed for the 3-back task (r = 0.252, p = 0.0415; Fig. 3I). Because no significant posterior cluster was observed for the 1-back task, this correlation was calculated using the temporal overlap of the significant 2-back and 3-back windows, 246–434 ms. No significant association was observed for the 1-back task (r = 0.071, p = 0.5724; Fig. 3G). Together, these findings indicate that trough rTMS improved working memory in parallel with stronger posterior task-related activity, and that the electrophysiological changes were related to the magnitude of behavioral improvement.

### rTMS enhances low-frequency network connectivity

Building on the behavioral effects and localized changes observed in task-related activity, we next investigated how rTMS affected broader inter-regional communication. Since local stimulation effects affects both local and connected, distributed brain networks (Dowdle et al., 2018; Güth et al., 2026; Wang et al., 2024), synchronized changes of oscillatory activity at the stimulated region and other areas can be measured as a consistent phase lag between the EEG channel nearest the stimulation site and the remaining channels. Hence, to investigate TMS-related phase-based connectivity changes, the debiased weighted phase lag index (dwPLI) was calculated between the EEG channel closest to the TMS coil (F3) and all remaining channels. The dwPLI reflects the degree of asymmetry between two phase distributions at two EEG channels. Using this method, our previous research in nonhuman primates demonstrated low-frequency connectivity changes to the stimulation site that scaled with TMS intensity in the 1-13 Hz range (Güth et al., 2026). This aligns with existing research highlighting the importance of low-frequency oscillations for long-range connectivity (Buzsáki & Draguhn, 2004). Consistent with this literature, the 5-second post-burst connectivity average across all subjects during rTMS exhibited the most notable increase in connectivity in the range of 1-4 Hz during the first second after bursts (Fig. 4A). Therefore, our analyses focused on this time-frequency range.

**Figure 4.**
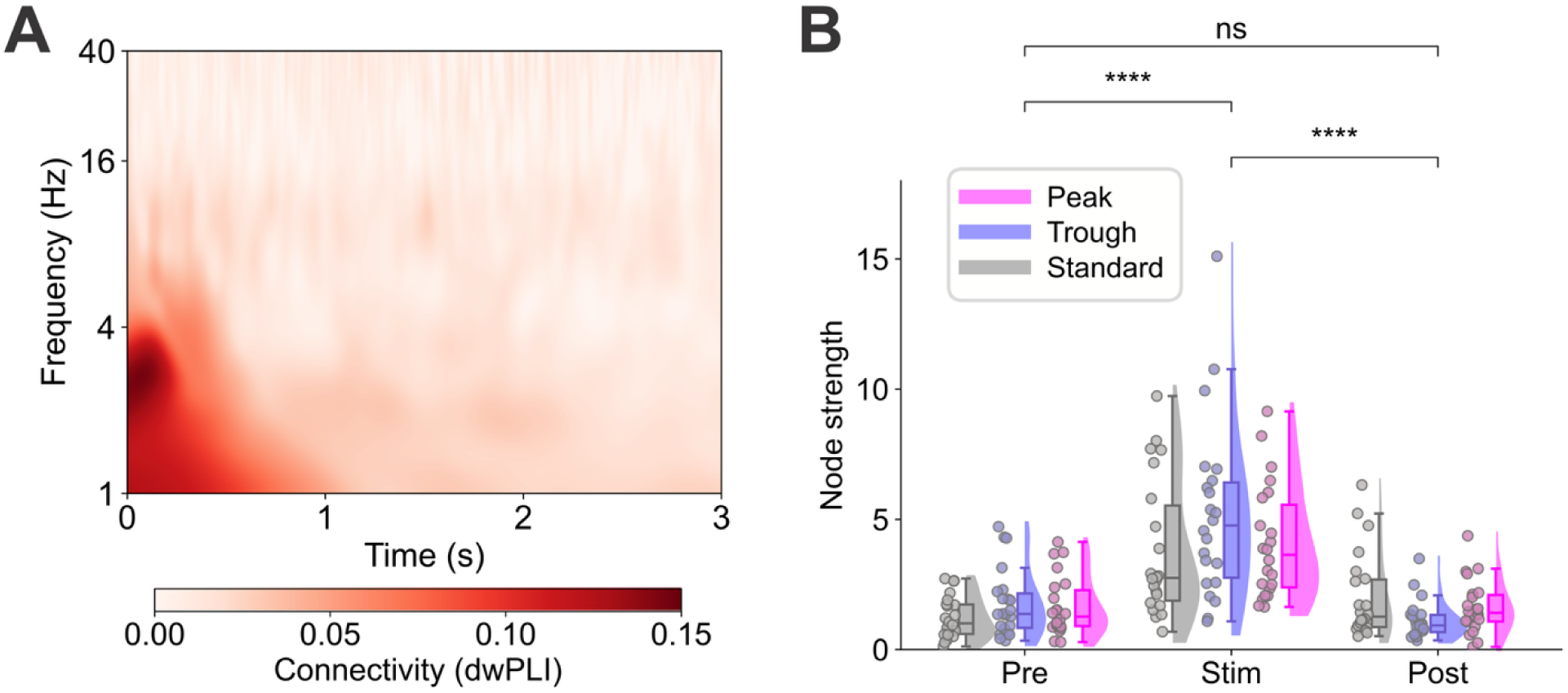
rTMS-induced changes in connectivity strength. A) Grand average of connectivity (debiased weighte phase lag index (dwPLI)) across all subjects and all channels during the stimulation block time-locked to the end of beta bursts. Data was convolved using a complex Morlet wavelet (3 cycles, 40 log-scaled frequencies from 1-40 Hz). B) Distributions of node strength for each group (Peak, Trough, Standard) and block (Pre: pre-stimulation, Stim: stimulation, Post: post-stimulation) calculated from the dwPLI values in the delta-theta range (1-4 Hz) over one second post-burst identified in A). ns: not significant, ****: p < 0.0001

To assess connectivity changes due to stimulation, beta-burst-induced connectivity was compared to 3-minute resting-state recordings pre- and post-stimulation. Graph theory was used to summarize connectivity patterns as networks of nodes (EEG channels) and edges (functional connections). For each participant and block (pre, stimulation, post), undirected weighted adjacency matrices were constructed to calculate node degree (number of connections above 95th percentile) and node strength (sum of all connection weights), reflecting the network size and overall connection strength respectively. Fig. 4B depicts the significant fixed effect of block from a linear mixed effects model predicting node strength (χ² = 25.36, p = 3.117e-06). Bonferroni-holm-corrected post-hoc comparisons showed that node strength significantly increased from the pre-stimulation to the stimulation block (β = 2.82, SE = 0.32, t(126) = 8.7, p_Holm_ = 5.458e-14) and decreased again in the post-stimulation block (β = -2.78, SE = 0.32, t(126) = -8.6, p_Holm_ = 4.527e-14). No significant changes from pre- to post-stimulation were found (β = 0.04, SE = 0.32, t(126) = 0.11, p_Holm_ = 0.914), and neither the fixed effect of group (χ² = 2.56, p = 0.278) nor its interaction with block reached significance (χ² = 9.4, p = 0.052).

### Phase-dependent anteroposterior network expansions

Figure 5A displays the most common F3-centered networks, illustrating the spatial extent of the stimulation induced networks across all groups. The most common connections were to left fronto-central and centro-parietal channels (e.g., C1) and along the midline (e.g., Fz). In addition to a significant main effect of block on node degree (χ² = 14.43, p = 0.001), a significant interaction of group and block revealed that changes of the network size across blocks differed by group (Fig. 5B; χ² = 15.35, p = 0.004). Only in the peak and trough groups, the F3-centered network significantly expanded from the pre-stimulation to the stimulation block (peak: β = 4.68, SE = 1.55, t(126) = 3.01, p_Holm_ = 0.006; trough: β = 5.86, SE = 1.55, t(126) = 3.77, p_Holm_ = 0.005). There was no such increase in the standard group (β = 2.64, SE = 1.55, t(126) = 1.7, p_Holm_ = 0.277). To explore which new connections were added through stimulation, Fig. 5C depicts the connections most commonly added during the stimulation block that were not present during the pre-stimulation block. In the peak group, the F3-centered network expanded to include centro-parietal (CP1, C1, Cz) and prefrontal (Fz, F1, FC3) connections in on average 50% of subjects. By contrast, network expansion in the trough group was spatially less consistent. With a mean of 44% of subjects, local fronto-central and left fronto-lateral channels (Fz, FC1, Af1, AF7), as well as two left centro-lateral channels (C3, T7) were added to the network. To verify the orientations of these spatial patterns we calculated the distributions of approximate signed anteroposterior distances to F3 based on the standard 10-10 electrode system (Fig. 5D). A significant Kruskal-Wallis test confirmed different spatial patterns between the peak, trough, and standard groups (H = 11.28, p = 0.004). Specifically, post-hoc Mann-Whitney U tests showed that the peak had a more posterior (U = 86, p_Holm_ = 0.004) and trough group a more anterior network distribution of expansions than the standard group (U = 116, p_Holm_ = 0.031). After the rTMS intervention, node degree significantly declined compared to the stimulation block in the peak (β = -5.46, SE = 1.55, t(126) = -3.51, p_Holm_ = 0.002) and trough groups (β = -9.86, SE = 1.55, t(126) = -6.35, p_Holm_ = 1.082e-08) but not in the standard group (β = -1.36, SE = 1.55, t(126) = -0.88, p_Holm_ = 0.764). In most peak and trough group subjects, the same connections that emerged during stimulation faded during the post-stimulation block (Supplementary Fig. 11). Finally, node degree decreased significantly from the pre-stimulation to the post-stimulation block only in the trough group (β = -4, SE = 1.55, t(126) = -2.57, p_Holm_ = 0.011). There was no clear spatial pattern as to which connections disappeared from the pre- to post-stimulation block across trough group subjects (Supplementary Fig. 12).

**Figure 5.**
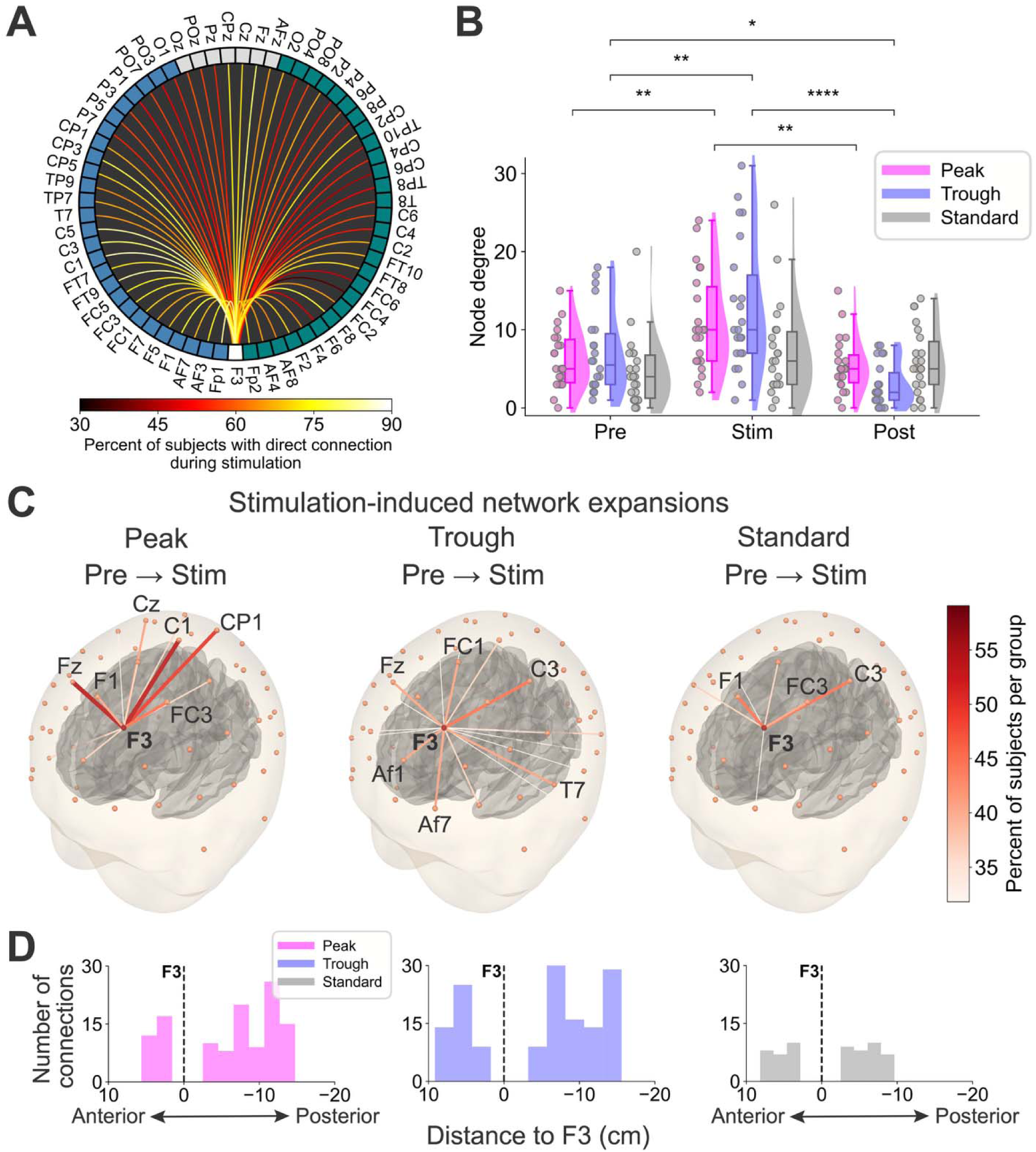
rTMS-induced network expansions. A) Circular connectivity plot showing the percent of subjects across all groups who had a given connection in their F3-centered network during the stimulation block. All connections ar relative to F3 (white node). Blue nodes highlight channels on the left hemisphere and teal nodes channels on the right hemisphere. Gray nodes mark the midline channels. B) Distributions of node degree for each group and block (Pre: pre-stimulation, Stim: stimulation, Post: post-stimulation) calculated from delta range (1-4 Hz) dwPLI values. C) New connections added to the network centered around F3 from the pre-stimulation block to the stimulation block separated by group. All three head models are based on the fsaverage template and show the electrode locations (orange-red spheres) from the montage used in this study on top of the scalp and the gray matter surface below. Th color and line thickness of each line represent how many subjects had this connection added to their network due t stimulation. Labels mark the name and location F3 and the most commonly added channel connection for a give group. D) Histograms of connections shown in C over signed anteroposterior distances to F3. Colors denote th group. Positive values indicate connections anterior of F3, negative values indicate connections posterior to F3. *: p < 0.05, **: p < 0.01, ****: p < 0.0001

## Discussion

This study tested whether the cognitive and neural effects of prefrontal beta-burst stimulation depend on the phase of ongoing beta activity. Using closed-loop rTMS targeting the left DLPFC, we found that phase-timed stimulation shaped executive performance across two cognitive domains: stimulation timed to the beta peak improved inhibitory-control performance during stimulation, whereas stimulation timed to the beta trough was associated with improved working-memory performance from pre- to post-stimulation. These behavioral effects were paralleled by condition-dependent changes in task-evoked ERPs, most prominently within the P3 time range, suggesting that stimulation altered post-stimulus processes related to inhibitory control and working-memory updating. Importantly, individual differences in stimulation-related behavioral change were associated with corresponding ERP changes, linking the cognitive effects of stimulation to task-related neural processing. Finally, phase-timed stimulation transiently altered DLPFC-centered connectivity, inducing anterior network expansion at the trough and posterior expansions at the peak. This suggests that ongoing beta phase shapes the susceptibility of prefrontal networks to stimulation. Together, these findings support the view that prefrontal beta phase indexes functionally relevant network states that can be leveraged to modulate executive function.

Specifically, the phase-function mapping that was observed in behavioral outcomes was corroborated by the physiological effects in the ERPs. Consistent with the reduction in commission errors, we observed increased fronto-central activity around 400 ms after No-go stimulus presentation, corresponding to the P3 component, when peak beta rTMS was applied. These observations align with the view that prefrontal beta activity and the No-go P3 are both involved in inhibitory control, while further suggesting that the phase of the ongoing beta rhythm shapes how this process is engaged. Indeed, absolute amplitude of the P3 was related to accuracy. Similarly, for working memory an increase in posterior midline components were observed between approximately 300 and 500 ms after stimulus presentation. This increase in amplitude was limited to trough beta stimulation, and was absent in peak beta and standard rTMS. Because P3 activity has been linked to updating, evaluation, and allocation of attentional resources, these findings suggest that trough-related beta stimulation affected working-memory performance by modulating post-stimulus processing of task-relevant information. Indeed, larger pre-to-post changes in working memory accuracy were associated with larger increases in posterior midline activity.

Our finding that stimulation effects depended on the phase of ongoing beta activity extends previous work on phase-targeted brain stimulation from cortical excitability and plasticity toward prefrontal cognition. Prior studies have shown that EEG-informed TMS can reveal phase-dependent differences in cortical excitability (Bergmann et al., 2012; Schaworonkow et al., 2019; Zrenner et al., 2018), particularly in the sensorimotor system, where stimulation timed to ongoing mu and beta rhythms has been used to probe state-dependent corticospinal responses (Haigh et al., 2025; Wischnewski et al., 2022). In repetitive protocols, phase-targeted stimulation has also been shown to shape after-effects of rTMS, suggesting that oscillatory phase influences whether short-term neural plasticity is induced (Faller et al., 2022; George et al., 2023; Zrenner et al., 2018). Initial evidence for phase-targeted stimulation of the prefrontal cortex was provided by Gordon et al. (2022), who showed that theta-phase-synchronized gamma-burst stimulation over the left dorsomedial prefrontal cortex produced phase-dependent changes in TMS-evoked prefrontal activity and theta-gamma coupling, alongside faster working-memory responses. More broadly, these findings suggest that the cognitive effects of prefrontal stimulation may depend on the combination of stimulation site, oscillatory frequency, and endogenous phase.

Phase-dependent stimulation effects may be especially informative in the DLPFC, where the same region contributes to multiple executive operations. The DLPFC is commonly viewed as a flexible control hub that supports the maintenance, selection, and suppression of task-relevant information, rather than a region dedicated to a single cognitive function (Cieslik et al., 2013; Friedman & Robbins, 2022; E. K. Miller & Cohen, 2001). This functional diversity raises the question of how activity within the same cortical region is organized to support different executive demands. Our findings hint at beta phase providing one mechanism for this flexibility. The pattern of behavioral effects is consistent with the possibility that inhibitory-control processes are preferentially linked to the peak of the prefrontal beta rhythm, whereas working-memory processes are preferentially linked to the trough. Therefore, our results suggest a potential beta phase mechanism for managing the diverse cognitive functions in the DLPFC by gating modes and networks of executive processing through its peak and trough phase.

In this view, phase may help determine which information is stabilized, routed, or suppressed within DLPFC-centered networks in addition to modulating excitability. Consistent with this notion, subjects in the trough group exhibited reduced network size post-compared to pre-stimulation, with no clear spatial trend as to which connections were removed. Homeostatic structural plasticity provides an interesting framework for speculating about both this stimulation-induced network reduction and the phase-dependent direction of network reorganization. Recent computational network modeling and in vitro tissue imaging combined with simulations demonstrated that network rewiring is driven by the system’s need to maintain functional equilibrium during ongoing perturbation (Anil et al., 2023; Lu et al., 2025). Specifically, structural plasticity operates under a biphasic rule, where the direction of network reorganization (i.e., synapse growth or pruning) depends on the degree of network disruption (Lu et al., 2025). Applying rTMS at a consistent phase (i.e., peak or trough) may rhythmically disrupt local DLPFC activity and its long-range communication mediated by low-frequency oscillations. Rather than causing a network failure, the phase-dependent disruption may force a homeostatic response, driving the network to reorganize in a more consistent fashion than standard rTMS. This may involve a lasting non-specific pruning of low-weight or unstable connections to optimize the reinforced network reorganization (Pascual-Leone et al., 2005).

In line with this interpretation, networks reorganized in distinct spatial orientations. In the peak group, network size increased with a posterior orientation, suggesting heightened communication within potentially frontoparietal and sensorimotor regions. This aligns with the peak group’s improvement in response inhibition, a process that critically relies on synchronized activity between prefrontal and motor regions (Aron et al., 2014; Swann et al., 2012). Conversely, in the trough group, network size increased with an anterior orientation, including EEG channels located above lateral as well as medial prefrontal regions. Coordinated interactions between these prefrontal subregions are commonly proposed as core mechanisms of working memory (Braver, 2012; Braver et al., 2007; Curtis & D’Esposito, 2003), which is consistent with the trough group’s improvement in working memory performance. However, despite this plausible alignment of spatial networks and gain in cognitive function, these observations remain speculative and are based only on EEG sensor locations. Further studies are needed to determine whether these connectivity shifts facilitate the observed behavioral changes or whether they represent unrelated physiological responses.

Beyond this limitation, inhibitory control was assessed during stimulation pauses, whereas working memory was assessed from pre-to post-stimulation. As a result, the phase-function mapping should be interpreted cautiously and may depend on the specific stimulation protocol and task context. Given the difference in timing of measuring working memory and inhibitory control in relation to the stimulation, the results of both effects cannot be directly compared. However, together they do provide evidence for phase-dependent processing of cognitive information. Future studies should aim at replicating the peak-inhibitory-trough-memory relationship that is observed here using comparable designs. Furthermore, it is important to investigate whether these observations transfer to other cognitive tasks, modalities, brain regions, and oscillatory rhythms. Although the precise phase dynamics underlying these effects require further verification, the present findings show that phase-targeted rTMS can produce behavioral and neural effects that are not observed under standard rTMS. This has important implications because rTMS is widely used as a therapeutic intervention for psychiatric and neurological disorders. Phase-targeted stimulation protocols may therefore offer a path toward improving the specificity and efficacy of clinical rTMS applications.

## Methods

### Participants

A sample of N=66 was collected (mean age ± SD: 22.39 ± 4.62, F/M ratio: 0.53/0.47). Before participation, volunteers were tested for eligibility based on the following inclusion criteria: I) Their age was between 18 and 45; II) there was no history of neurological or psychiatric disorders; III) participants had no history of seizures or epilepsy, nor did they have direct family members with epilepsy; IV) participants had no recent head injuries or metallic objects in head and neck area; V) participants understood English and were able and willing to give informed consent. Approval for this study was given by the institutional review board of the University of Minnesota, and all procedures followed the declaration of Helsinki.

Sample size was based on a power analysis using G*Power, V3.1 (Faul et al., 2009). Assuming a between-subjects ANOVA, alpha = 0.05, power (1-beta) = 0.95, and a moderate effect size of f = 0.25, the analysis suggested a sample size of N=66.

### Electroencephalography

EEG data was recorded with a 64 active channel (according to the international 10-20 system) EEG system (actiCAP EEG cap, actiCHamp amplifier; Brain Products GmbH, Gilching, Germany). Data was recorded on Matlab via LabStreaming Layer (during phase-specific rTMS) and via BrainVision Recorder (before and after rTMS). The sampling rate in all instances was 1000 Hz and saved for offline analysis. Electrode impedances were kept below 10 kΩ. EEG was recorded for a 4-minute resting-state (eyes open) and the working memory task (explained below) before and after rTMS, as well as during rTMS for phase targeting and the Go/Nogo task (explained below).

### Oscillation phase algorithm

In this experiment, stimulation bursts were applied at distinct phases of the prefrontal beta rhythm. To that end, neural oscillations were tracked online with EEG. The EEG data was streamed to Matlab 2023b online using Lab Streaming Layer (github: https://github.com/sccn/labstreaminglayer, Kothe et al., 2025), with <1 ms precision. The Bayesian Temporal Prediction (BTP) algorithm was used to identify and predict EEG phase in the live data stream (Shirinpour et al., 2025). To optimize the parameters used for real-time phase prediction and improve the accuracy of online pulse delivery, the BTP algorithm was first trained on a 3-minute resting state EEG period prior to the experiment, as described in Shirinpour et al. (2025). During the real-time EEG analysis, the system identifies a reliable anchor point within the filtered signal and projects forward to the target phase using prediction parameters optimized for each subject (e.g., window length, projection distance), providing robust real time phase estimation for closed loop stimulation. The sampling rate was 1000 Hz, and online data for phase prediction was extracted from electrode position F3 (according to the international 10-20 system). The data was spatially filtered by applying a Laplacian reference using eight surrounding electrodes (Fp1, AFz, AF7, Fz, F7, FC1, FC3, FC5).

### Phase-specific transcranial magnetic stimulation

The individualized TMS paradigm used a modified version of a standard 20 Hz protocol, which is used for treatment of depression and obsessive compulsive disorder (Cohen et al., 2022). In the standard 20 Hz paradigm, which here was applied as a control condition, a burst of 40 pulses is applied in 2 seconds with 50 ms between pulses. Between each burst there is a period of 20 seconds without stimulation. In this control condition (standard TMS) oscillation phase is not considered and therefore TMS is applied at random phases of the beta oscillation. It is important to emphasize that we consider standard TMS as a control condition in this experiment to isolate the effects of phase-dependent stimulation. Nevertheless, 20 Hz rTMS is commonly used as an active intervention in research and clinical contexts. In the two phase-specific TMS conditions also bursts of 40 pulses are applied. However, the time between pulses depended on each participant’s individualized inter-cycle interval of beta oscillations, which was estimated from the resting state EEG when the BTP algorithm was trained. In the peak target condition, the peak of the beta oscillation is targeted with the first pulse, followed by the remaining 39 pulses with an inter-stimulus interval that is equivalent to the beta inter-cycle interval. The trough condition was similar, with the difference that the first pulse of the burst was locked to the trough of the beta oscillation. Note that TMS significantly disrupts ongoing EEG, and therefore, it was not feasible to phase-target each individual TMS pulse.

In all experimental conditions, the TMS coil was placed over the left DLPFC, at MNI coordinates [-40, 34, 28] and at an angle of 45° to the midline. This region was based on meta-analytic evidence showing the involvement of this region in inhibitory control (Carlisi et al., 2017). In the international 10-20 system, these MNI coordinates are closest to electrode F3, which was used for beta signal extraction. The TMS coil was positioned and kept in place at this location with a Brainsight neuronavigation system (Rogue Research Inc., Montreal, Canada) and using the standard 152-MNI head model. The coil was held by the experimenter, which has an advantage compared a rigid mechanical arm, as movements of the participant can be controlled for. Stimulation was always applied at the target location within a margin of 2 mm and an angle deviation of 2°.

The intensity of stimulation was set to 120% of the resting motor threshold (rMT). The rMT was determined by placing the TMS coil over the primary motor cortex hotspot corresponding to the hotspot of the right first dorsal interroseus hand muscle at an angle of 45° to the midline. First, the hotspot is determined by finding the largest response in a quasi-random grid of the motor cortex. Subsequently, an adaptive threshold-hunting algorithm is used to determine the minimal intensity at which a muscle response is observed. This intensity is multiplied by 1.2 and used for DLPFC stimulation.

### Cognitive tasks

To measure response inhibition, a go/no-go task was used, which was programmed in MATLAB 2023b. In this task, participants are instructed to withhold a response for a particular stimulus, while in the majority of trials, they need to press a key on a Cedrus response box. Specifically, participants were shown either a square, a circle, or a triangle in a ratio of 4:1:1. Both the square and circle represented go stimuli, with the former a common and the latter a rare go stimulus. The addition of a rare go stimulus increases the difficulty of the task, which was used to avoid ceiling effects in accuracy. The triangle was the no-go stimulus where participants had to withhold their response. Stimuli were shown for 100 ms, followed by a fixation cross, which was presented for a jittered duration between 700 and 1000 ms. This task was interleaved with the stimulation paradigm, meaning that 10 trials were presented in the 25 seconds between stimulation bursts. This was repeated for 48 blocks, resulting in a total of 480 trials (320 common go, 80 rare go, and 80 no-go trials). The number of commission errors and reaction time were used as outcome measures for the go/no-go task.

To measure working memory, a verbal n-back task was used with three levels of difficulty, and was programmed in MATLAB 2023b. The three levels were a 1-back, 2-back, or 3-back task. Stimuli consisted of eight consonants (C, F, K, N, S, T, V, and Z). Each stimulus was displayed for 250 ms, followed by a fixation cross, which was displayed for a jittered interval between 1250 and 1750 ms. After each letter, the participants had to indicate whether the letter had appeared before on a Cedrus response box (green button: yes, red button: no). For the 1-back, participants indicated whether the current letter was the same as the previous letter. For the 2-back task, they indicated whether the current letter was the same as the letter two trials ago. For the 3-back task, they indicated whether the current letter was the same as the letter three trials ago. Each n-back level consisted of 120 trials, with a n-back chance of 25%. D-prime (D’ = Z_hit_rate_ - Z_false_alarms_) and reaction time were used as outcome measures for the 1-, 2-, and 3-back.

### Event-related potential preprocessing

Event-related potentials (ERPs) were computed in MATLAB and the FieldTrip toolbox (Oostenveld et al., 2011). Continuous EEG data were epoched from −500 to +1000 ms relative to stimulus onset and preprocessed prior to averaging. The data was re-referenced to an average reference followed by detrending and demeaning using a baseline correction window from −200 to 0 ms. Data were band-pass filtered between 1 and 50 Hz using a two-pass Butterworth filter as implemented in FieldTrip. An independent component analysis was applied to identify artifacts. Components were classified as artifact-related based on their scalp distribution and time course, with particular attention to components reflecting eye blinks, eye movements, or other stereotyped non-neural activity. Artifact-related components were removed from the data. Remaining noisy trials were rejected using visual inspection with FieldTrip’s summary-based rejection procedure. On average, 2.73 ± 1.18 ICA components were removed for the Go/No-Go task and 3.51 ± 1.78 components for the N-back task. Visual trial rejection removed 4.95 ± 4.07% of trials in the Go/No-Go task and 6.12 ± 4.89% of trials across N-back task sessions; detailed values are provided in Supplementary Table 1.

For the Go/No-Go task, ERPs were time-locked to NoGo trials. For the N-back task ERPs were time-locked to letter presentation. For both tasks, participant-level ERPs were aggregated into stimulation groups (peak, trough, and standard stimulation), and grand-average ERPs were computed. Previously, it has been shown that cognitive components related to the Go/No-Go and N-back task center around frontal and posterior midline regions, as reflected by N200 and P300 components (Albert et al., 2013; Scharinger et al., 2017). Therefore, for statistical analyses two regions of interest (ROI) were defined: A frontal ROI, taking the average of channels F1, Fz, F2, FC1, FCz, FC2, and Cz, and a posterior ROI, taking the average of channels Cz, CPz, Pz, CP1, CP2, P1, and P2.

### Analysis of Behavioral Data

Behavioral analyses were performed separately for the Go/No-Go and N-back tasks. For the Go/No-Go task, behavioral performance was quantified as the number of commission errors and reaction times. Reaction times were analyzed in milliseconds and trials with reaction times below 100 ms or exceeding three standard deviations above the participant-specific mean were excluded from reaction-time analyses. For the N-back task, performance was quantified using d-prime (standardized difference between the hit rate and false-alarm rate).

For the Go/No-Go task one-way ANOVAs were performed with number of commission errors and reaction time as dependent variables. Stimulation condition (peak, trough, standard) was included as the between-subjects factor. For the N-back task, pre-to-post rTMS change scores were calculated by subtracting pre-stimulation performance from post-stimulation performance. These change scores were used as dependent variables in mixed ANOVAs with N-back level as a within-subject factor and stimulation condition as a between-subject factor. Significant effects were followed by post hoc tests, using separate one-way ANOVAs for each N-back level and independent-samples t-tests comparing peak, trough, and standard stimulation conditions. Post hoc p-values were Bonferroni-corrected (p_Bonf_). Partial eta squared (ηp²) and Cohen’s d was calculated as a measure of effect size.

### Event-related potential analysis

ERP data were analyzed using cluster-based permutation testing (Maris & Oostenveld, 2007). Analyses were performed separately for each task and ROI (frontal, posterior) using 5000 Monte Carlo permutations, with clustering restricted to the 100–600 ms post-stimulus interval. Cluster correction used a cluster-forming threshold of *p* < 0.05 and the maximum cluster-sum statistic. Spatial neighbours were defined from the electrode layout. For the Go/No-Go task, NoGo ERPs were compared between stimulation groups using a between-subjects cluster-based one-way ANOVA. For the N-back task, post-minus-pre ERP difference waves were computed separately for each participant, stimulation condition, and N-back level. These difference waves were then tested separately for the 1-back, 2-back, and 3-back tasks using between-subjects cluster-based one-way ANOVAs with stimulation group as the independent variable. Significant clusters were followed up by extracting the mean ERP amplitude within the cluster window and performing Bonferroni-corrected independent-samples t-tests between stimulation conditions. Cohen’s d was calculated for post hoc comparisons. For visualization, scalp topographies were generated by calculating the descriptive one-way ANOVA F-value for each channel, averaged across the significant cluster time window. These topographies were used only to illustrate the spatial distribution of the effect and were not used for additional statistical inference.

Brain–behavior associations were assessed using Pearson correlations pooled across stimulation conditions. For the Go/No-Go task, mean ERP amplitude in the significant cluster window was correlated with the number of commission errors. For the N-back task, ERP amplitudes in significant clusters were correlated with pre-to-post change in d-prime. Because no significant posterior 1-back cluster was observed, the 1-back correlation used the temporal overlap of the significant posterior 2-back and 3-back cluster windows.

### Connectivity preprocessing

EEG data recorded during the pre- and post-stimulation rest blocks as well as during the stimulation block were preprocessed in MNE-Python (Gramfort et al., 2013). Stimulation artifacts were linearly interpolated from -20 ms to the end of the burst duration. Next, continuous EEG data were band-pass filtered between 0.1 and 60 Hz using and an independent component analysis was run to identify artifact components. Interpolated stimulation periods were excluded from this to avoid biasing the decomposition. Analogously to the ERP preprocessing, components were classified as artifact-related and excluded from back-projection based on their scalp distribution and time course. On average 6.48 ± 4.4 components were removed for the pre-stimulation block, 6.4 ± 1.87 for the stimulation block, and 6.65 ± 4.33 for the post-stimulation block. Finally, continuous data were re-referenced to the average of all EEG channels and epoched from 0 to +5000 ms relative to the end of each burst. For the pre- and post-stimulation blocks, 48 overlapping 5-second epochs (matching the number of stimulation bursts) were extracted with randomly jittered onset times.

### Phase-based Connectivity

To investigate TMS-related connectivity changes, phase-based connectivity was calculated between all sensor pairs capturing potential network-wide changes using MNE-Python’s spectral_connectivity_epochs (Gramfort et al., 2013). TMS pulses delivered over the left DLPFC affects local neural activity and the activity of connected, distributed networks (Dowdle et al., 2018; Güth et al., 2026; Wang et al., 2024). Such propagation of synchronized, pulse-locked changes of oscillatory activity at the stimulated region and other areas can be measured as a consistent phase lag between the EEG channel nearest the stimulation site and the remaining channels (Güth et al., 2026). Therefore, connectivity pairs were masked to isolate pairs including the channel closest to the coil (F3). Connectivity values were calculated based on the discrete Fourier transform of the preprocessed post-stimulation epochs. Exploration of the mean connectivity changes averaged across all groups and burst epochs revealed the largest connectivity increase was in the range of 1-4 Hz from 0 to 1000 ms after the bursts. Hence, subsequent analyses focused on this range. Connectivity was calculated as the debiased weighted phase lag index (dwPLI) (Vinck et al., 2011). The dwPLI reflects the degree of asymmetry between two phase distributions at two EEG channels, where a dwPLI of 1 indicates a consistent phase-lag relationship and a dwPLI of 0 indicates no consistent phase lag. Thus, the dwPLI assesses connectivity as the consistency of synchronized oscillatory activity between two sensors. Importantly, zero-lag connections are excluded from the dwPLI, rendering it less susceptible to potential pulse-locked residual artifacts with near instant propagation. Compared to the conventional phase lag index, the dwPLI is also more robust to other confounding factors. First, it weights the contribution of observed phase leads and lags by the magnitude of the imaginary component of the cross-spectrum, reducing sensitivity to random noise. Second, it includes a correction for small sample sizes. To assess connectivity across sessions, beta burst-induced connectivity was compared to 3-minute resting-state recordings pre- and post-stimulation. Random overlapping segments were sampled from resting-state EEG and analyzed identically to the stimulation epochs (40 segments, 1-second windows).

### Graph Theory Measures

Graph theory was used to summarize connectivity patterns as networks of nodes (EEG channels) and edges (functional connections) using the networkx package in Python (Hagberg et al., 2008). For each participant and block (pre, stimulation, post), undirected weighted adjacency matrices were constructed from the dwPLI values. Negative values were set to zero. To focus on the most robust interactions, a proportional threshold was applied within each participant and block by retaining connections at or above the 95^th^ percentile of the connectivity matrix. The resulting matrix was binarized and used to construct an undirected graph. To evaluate TMS-induced network changes, node degree at F3 was calculated as the number of connections to F3 in the thresholded binary graph, reflecting changes of the network extent. Node strength was computed from the unthresholded weighted connectivity matrix as the sum of all edge weights connected to F3, reflecting overall connection magnitude changes due to TMS. All graphs were treated as undirected because the dwPLI is directionless, allowing us to account for both direct and indirect TMS-induced network effects. Differences in node strength and node degree between groups and blocks were tested using linear mixed effects models with fixed effects terms for group (peak, trough, standard) and for block (pre, stimulation, post), their interaction term, as well as a random intercept for each subject. Fixed effects were tested for significance using Wald tests, and post-hoc pairwise comparisons were performed on estimated marginal means. To account for the substantially higher number of post-hoc comparisons introduced by the addition of block as a fixed effect, p-values were corrected using the Bonferroni-Holm (p_Holm_) instead of the Bonferroni method.

### Directional network expansion analysis

To explore directional changes in network expansions from the pre-stimulation to the stimulation block, we identified newly formed connectivity edges and calculated their Euclidean distance to F3 (in cm) based on standard electrode coordinates for each subject. Distances were signed along the anterior-posterior axis (y-plane) to define whether locations were posterior (negative) or anterior to F3 (positive). To only retain robust connections, the sum of connections was calculated across subjects and only channel connections that occurred in over 30% of subjects were maintained. For each subject and group, the spatial distance to F3 was weighted by the counts of newly formed edges to derive a metric reflecting group-level directional network change. To test whether the three groups had different weighted mean directions of connectivity expansion relative to F3 a Kruskal-Wallis test was run. Post-hoc comparisons between groups were run using two-sided Mann-Whitney U tests. P-values were corrected using the Bonferroni-Holm method.

