## Supplementary Materials for "Phase-specific stimulation of prefrontal beta oscillations modulates distinct cognitive processes"

| Supplementary Table 1. EEG cleaning. Amount of removed ICA components and percentage or removed trials in the Go/Nogo and N-back task. | | | | |
| --- | --- | --- | --- | --- |
|  | Removed ICA components, mean | Removed ICA components, SD | Removed trials (%), mean | Removed trials (%), SD |
| Go/Nogo | 2.73 | 1.18 | 4.95 | 4.07 |
| 1-back pre | 3.74 | 2.04 | 5.54 | 4.64 |
| 1-back post | 3.44 | 2.03 | 6.19 | 4.70 |
| 2-back pre | 3.39 | 1.79 | 6.25 | 6.93 |
| 2-back post | 3.62 | 1.61 | 6.07 | 3.89 |
| 3-back pre | 3.45 | 1.51 | 6.15 | 4.70 |
| 3-back post | 3.41 | 1.69 | 6.53 | 3.97 |
| N-back total | 3.51 | 1.78 | 6.12 | 4.89 |

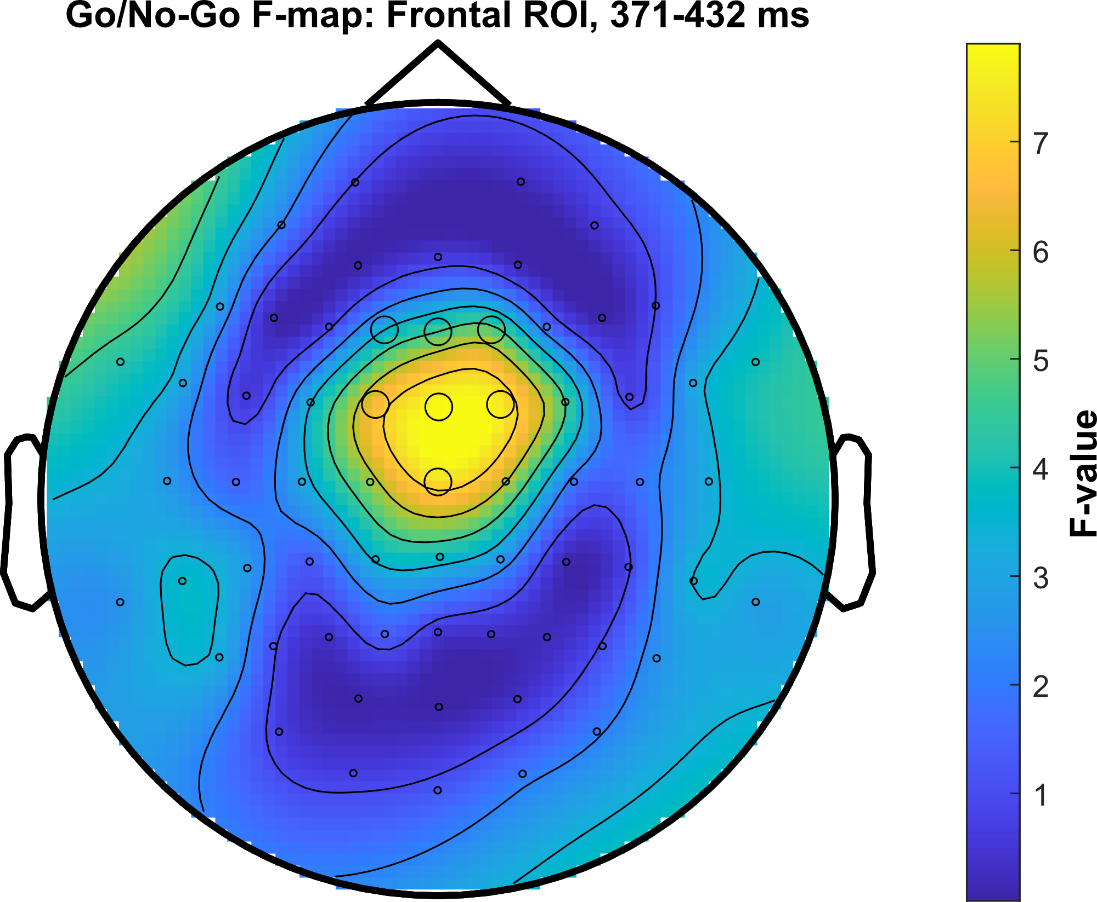

**Supplementary Figure 1. Scalp distribution of stimulation-dependent differences in NoGo event-related potentials.** Topographical representation of F-values obtained from a one-way between-subjects ANOVA comparing NoGo ERP amplitudes across the three stimulation conditions (peak-phase, trough-phase, and standard 20 Hz rTMS). F-values were calculated separately for each EEG electrode, averaged over the 371–432 ms post-stimulus interval, corresponding to the P3 component. The larger black circles indicate the five electrodes included in the predefined midline ROI (Fz, FCz, Cz, CPz, and Pz). This time window was identified using cluster-based permutation testing within the predefined frontal midline region of interest (ROI), revealing a significant effect of stimulation condition (FClus = 1965.75, p = 0.0378). Warmer colors indicate larger differences between stimulation conditions, with the strongest effects observed over frontocentral electrodes.

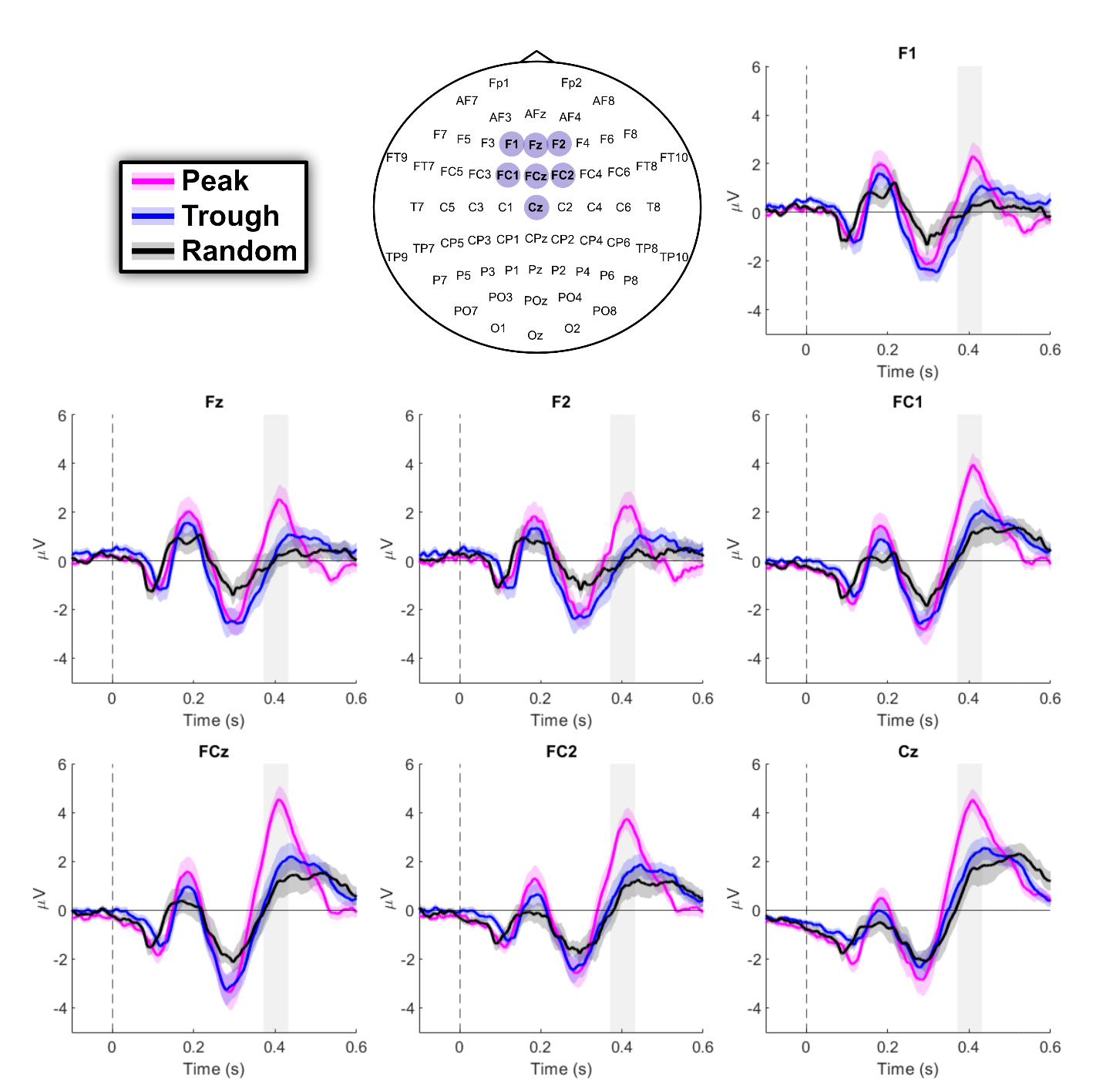

**Supplementary Figure 2. NoGo event-related potentials across frontocentral electrodes.** Grand-average ERP waveforms during the Go/NoGo task are shown for peak-phase (magenta), trough-phase (blue), and standard 20 Hz rTMS (black) across seven frontocentral electrodes (F1, Fz, F2, FC1, FCz, FC2, and Cz), highlighted in the electrode layout. The dashed vertical lines indicate stimulus onset (0 ms), while the grey shaded areas indicate the 371–432 ms time window in which significant differences between stimulation conditions were identified using cluster-based permutation testing. Across electrodes, peak-phase stimulation was associated with a larger positive ERP deflection during this time window. Shaded areas around each line represent the standard error of the mean across participants.

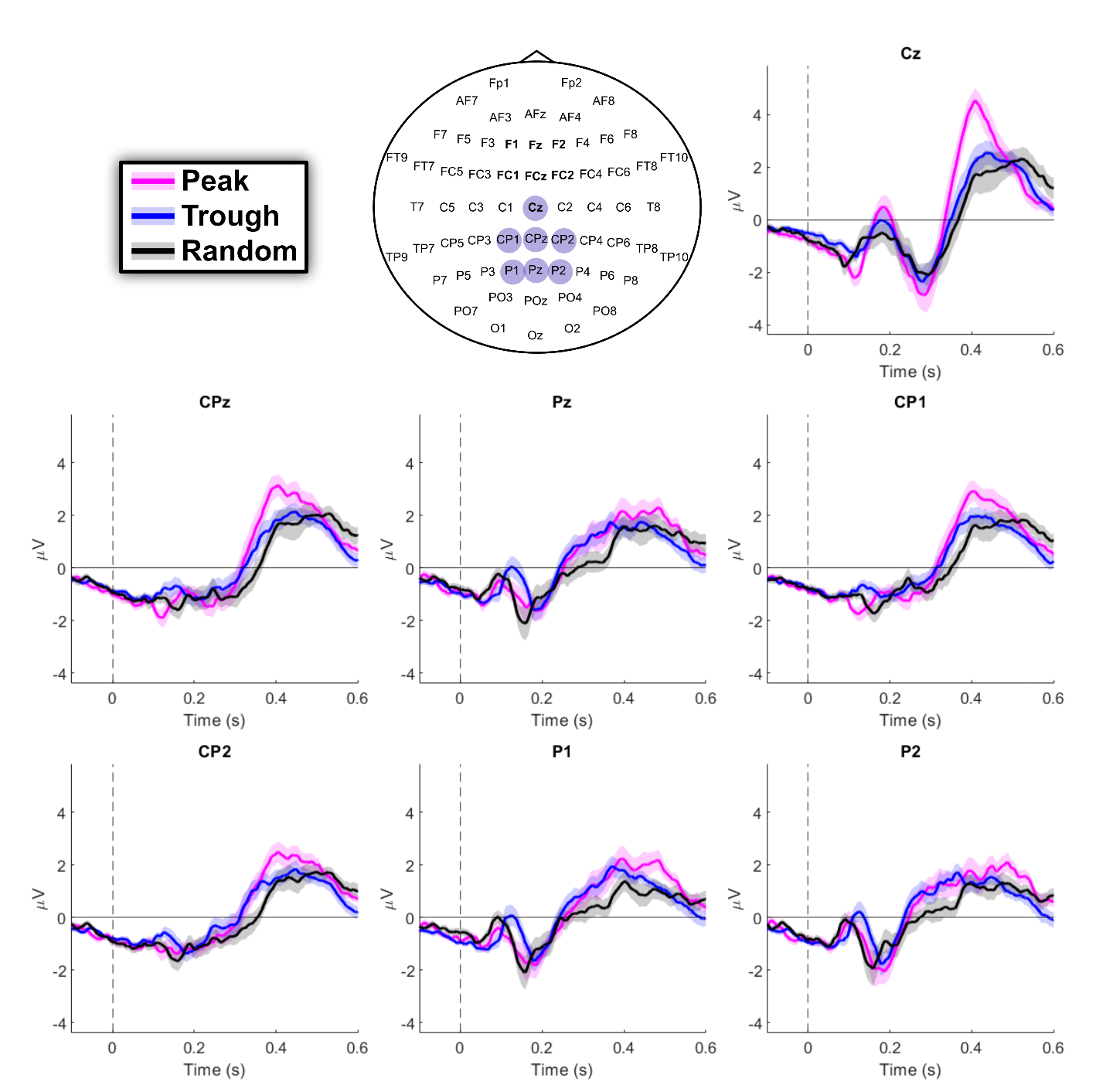

**Supplementary Figure 3. NoGo event-related potentials across posterior electrodes.** Grand-average ERP waveforms during the Go/NoGo task are shown for peak-phase (magenta), trough-phase (blue), and standard 20 Hz rTMS (black) across seven central and posterior electrodes (Cz, CPz, Pz, CP1, CP2, P1, and P2), highlighted in the electrode layout. The dashed vertical lines indicate stimulus onset (0 ms). Cluster-based permutation testing within the predefined posterior ROI revealed no significant differences between stimulation conditions. Shaded areas around each line represent the standard error of the mean across participants.

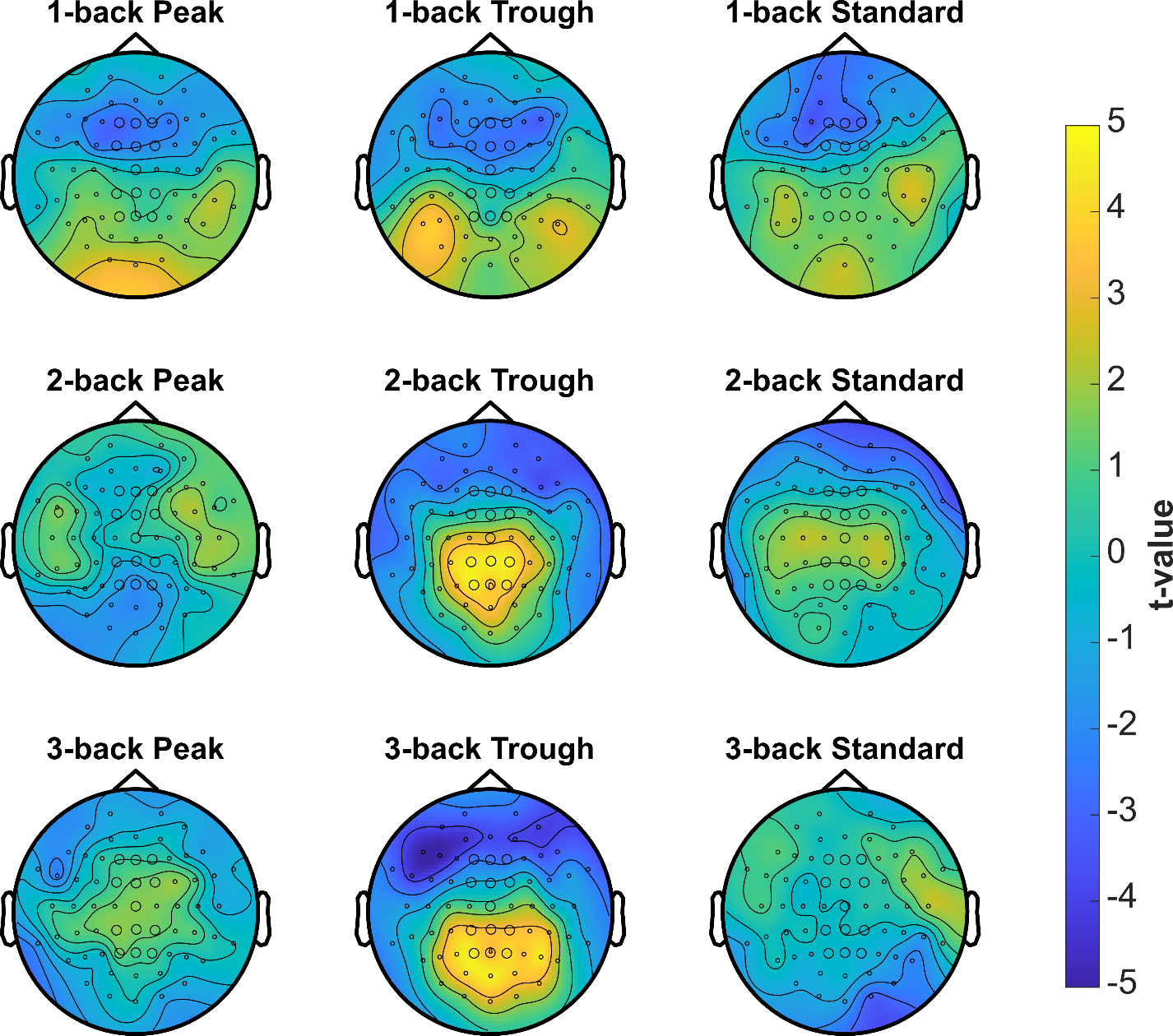

**Supplementary Figure 4. Scalp topographies of pre-to-post ERP changes across stimulation conditions and N-back levels.** Topographical maps show electrode-wise t-values for pre-to-post changes in ERP amplitude separately for each stimulation condition and N-back level. Rows correspond to the 1-back, 2-back, and 3-back tasks, and columns correspond to the peak-phase, trough-phase, and standard 20 Hz rTMS conditions. Warmer colors indicate more positive pre-to-post changes, whereas cooler colors indicate more negative pre-to-post changes. The larger black circles indicate the electrodes included in the predefined frontal and posterior regions of interest. These topographical maps are shown to illustrate the spatial distribution of ERP changes across the scalp and were not subjected to additional electrode-wise significance testing.

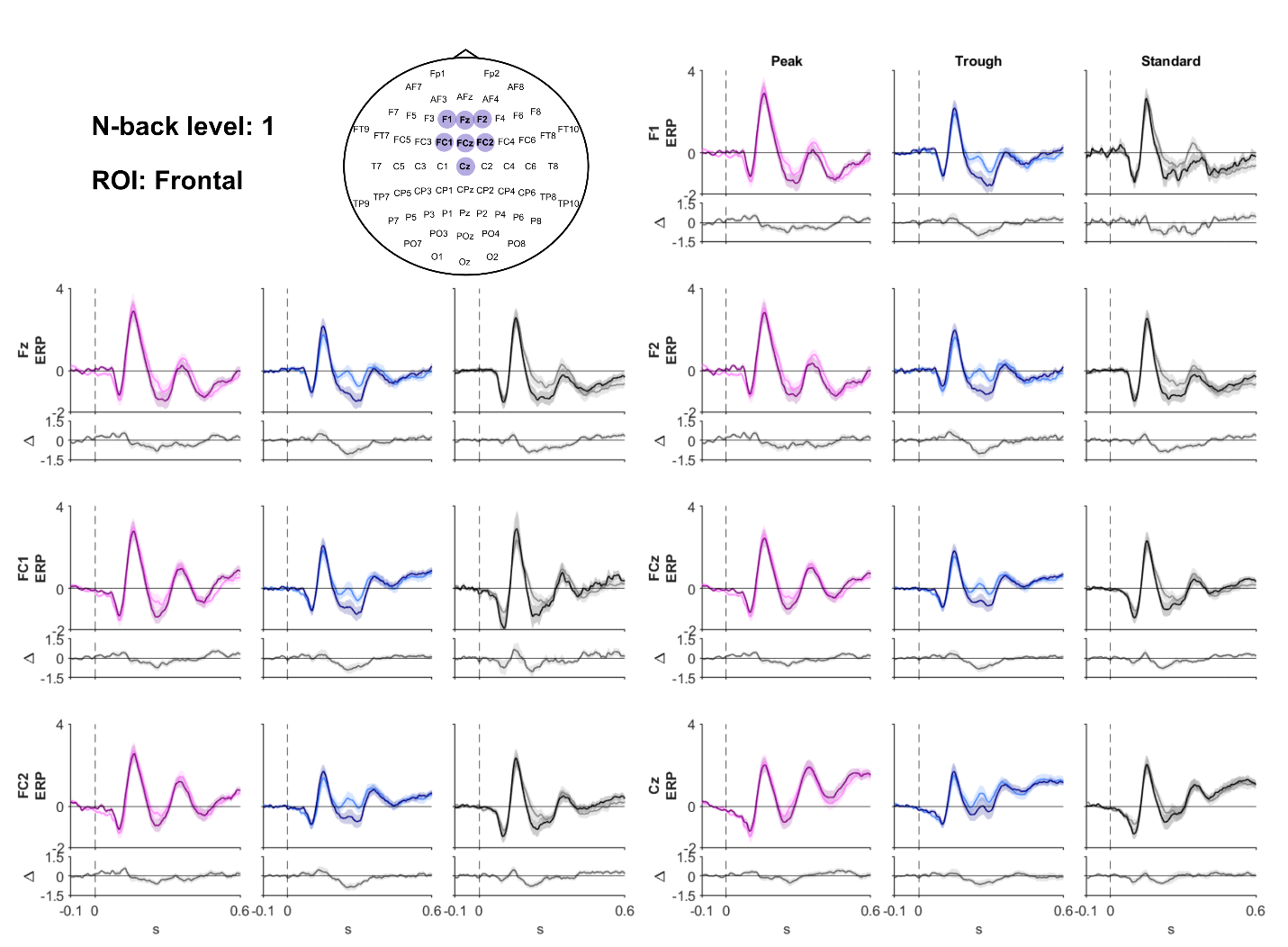

**Supplementary Figure 5. Event-related potentials across frontal electrodes during the 1-back task.** Grand-average ERP waveforms recorded before and after rTMS are shown for peak-phase (magenta), trough-phase (blue), and standard 20 Hz rTMS (black) across seven frontal and frontocentral electrodes (F1, Fz, F2, FC1, FCz, FC2, and Cz), highlighted in the electrode layout. The shaded bands surrounding the waveforms represent the standard error of the mean (SEM) across participants. The lower panels show the corresponding post-minus-pre stimulation difference waves (Δ) for each electrode and stimulation condition. Dashed vertical lines indicate stimulus onset (0 ms). Cluster-based permutation testing revealed no significant differences in pre-to-post ERP changes between stimulation conditions within the frontal region of interest.

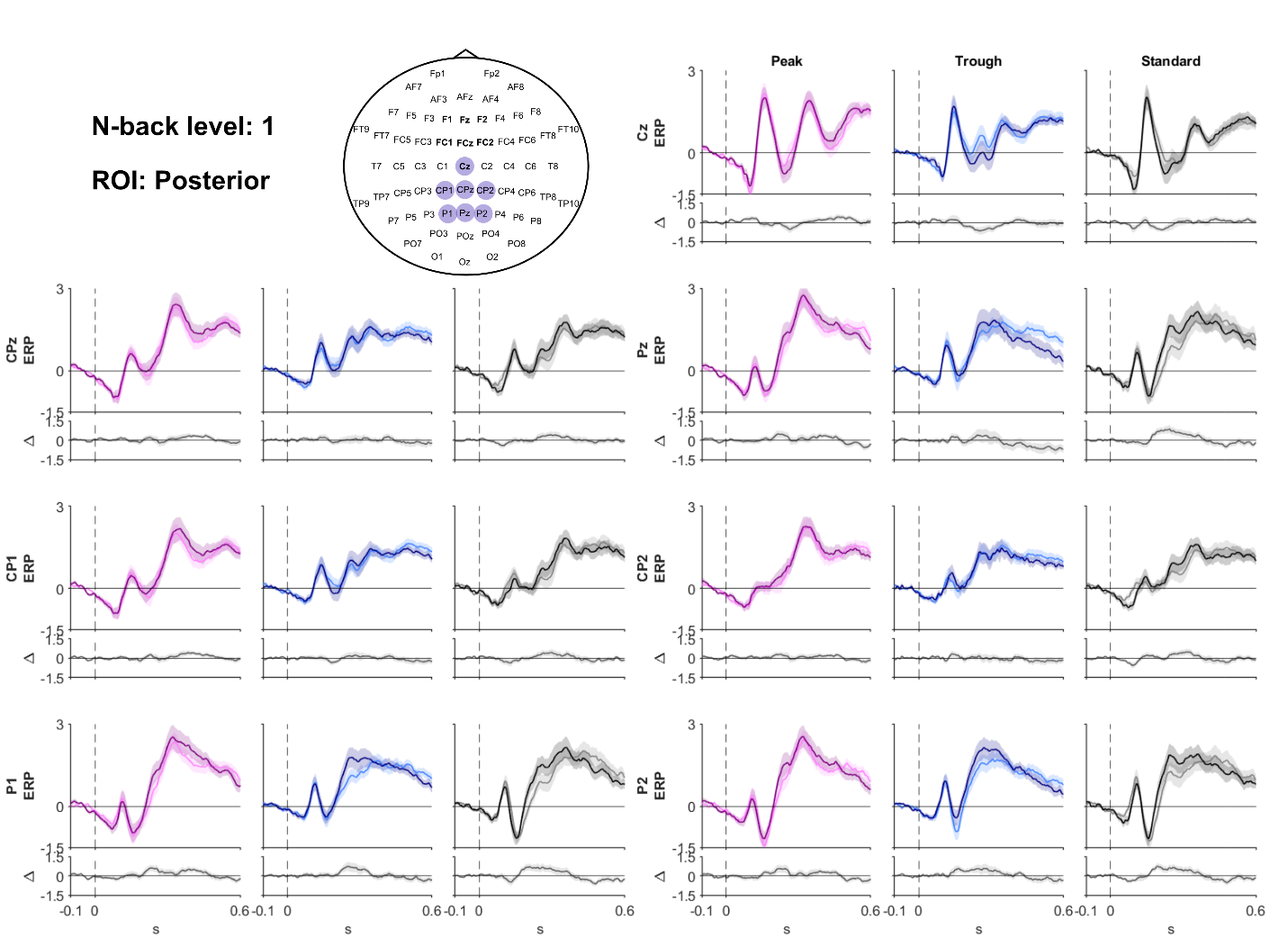

**Supplementary Figure 6. Event-related potentials across posterior electrodes during the 1-back task.** Grand-average ERP waveforms recorded before and after rTMS are shown for peak-phase (magenta), trough-phase (blue), and standard 20 Hz rTMS (black) across seven central and posterior electrodes (Cz, CPz, Pz, CP1, CP2, P1, and P2), highlighted in the electrode layout. The shaded bands surrounding the waveforms represent the standard error of the mean (SEM) across participants. The lower panels show the corresponding post-minus-pre stimulation difference waves (Δ) for each electrode and stimulation condition. Dashed vertical lines indicate stimulus onset (0 ms). Cluster-based permutation testing revealed no significant differences in pre-to-post ERP changes between stimulation conditions within the posterior region of interest.

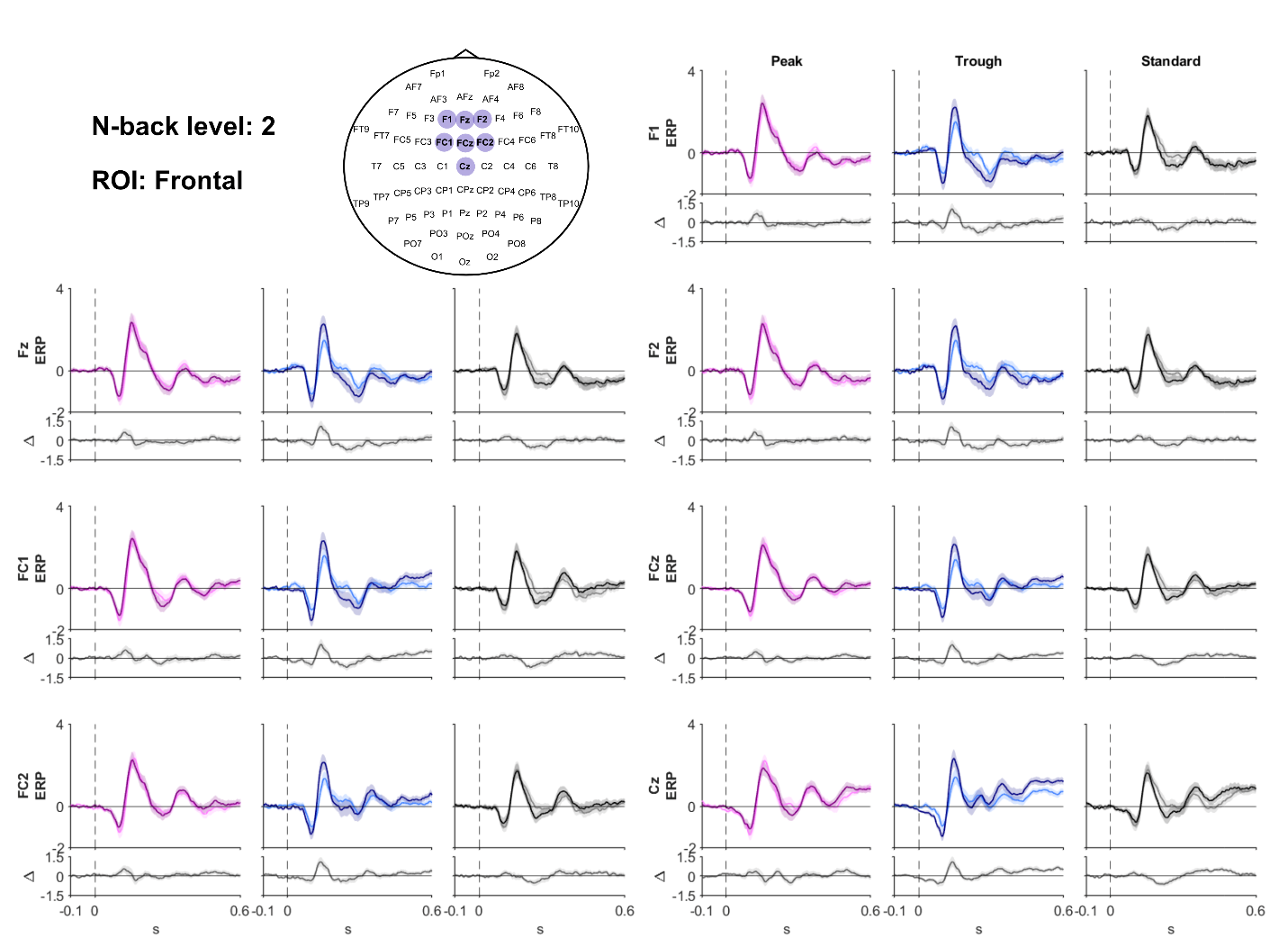

**Supplementary Figure 7. Event-related potentials across frontal electrodes during the 2-back task.** Grand-average ERP waveforms recorded before and after rTMS are shown for peak-phase (magenta), trough-phase (blue), and standard 20 Hz rTMS (black) across seven frontal and frontocentral electrodes (F1, Fz, F2, FC1, FCz, FC2, and Cz), highlighted in the electrode layout. The shaded bands surrounding the waveforms represent the standard error of the mean (SEM) across participants. The lower panels show the corresponding post-minus-pre stimulation difference waves (Δ) for each electrode and stimulation condition. Dashed vertical lines indicate stimulus onset (0 ms). Cluster-based permutation testing revealed no significant differences in pre-to-post ERP changes between stimulation conditions within the frontal region of interest.

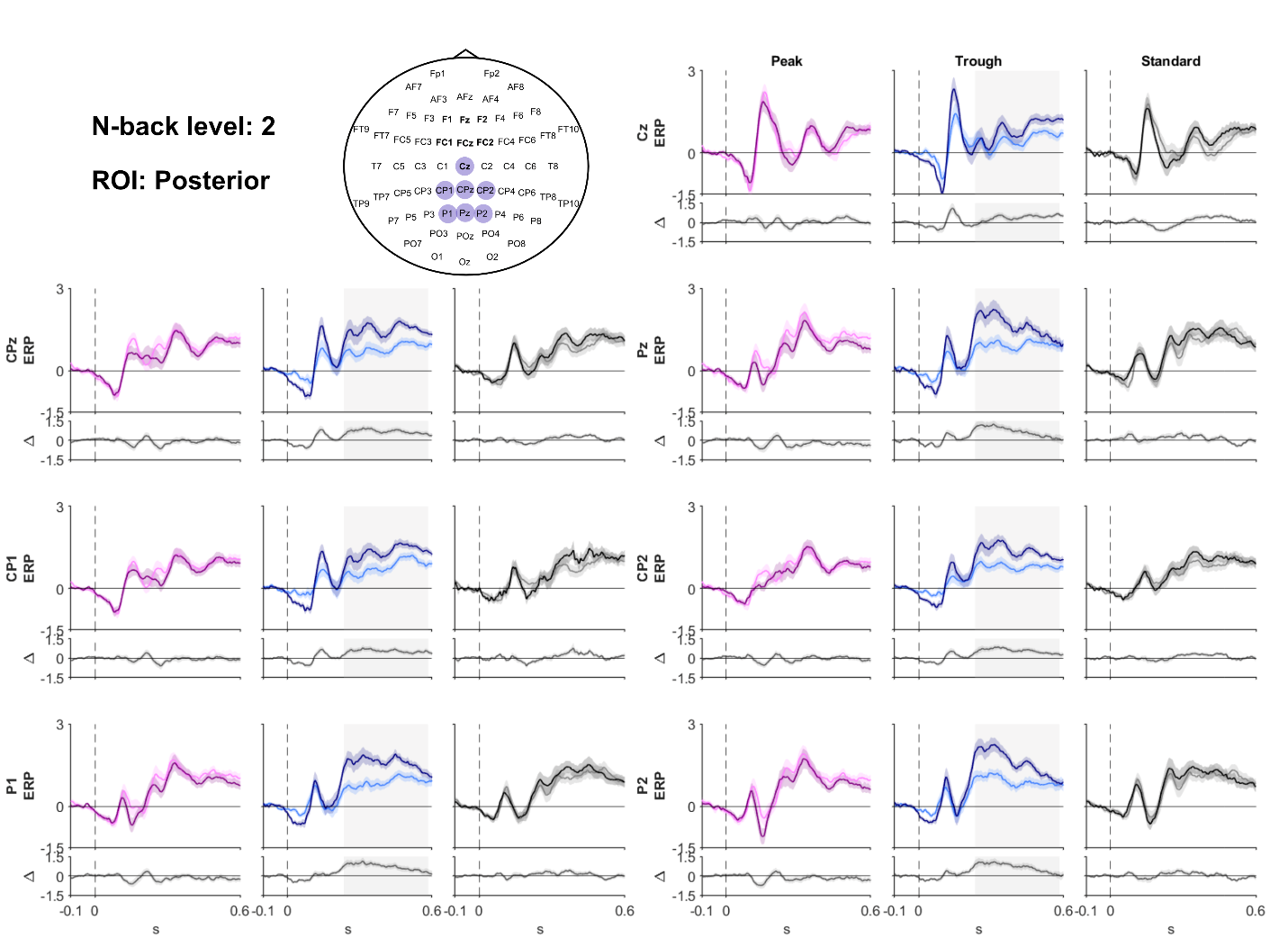

**Supplementary Figure 8. Event-related potentials across posterior electrodes during the 2-back task.** Grand-average ERP waveforms recorded before and after rTMS are shown for peak-phase (magenta), trough-phase (blue), and standard 20 Hz rTMS (black) across seven central and posterior electrodes (Cz, CPz, Pz, CP1, CP2, P1, and P2), highlighted in the electrode layout. The shaded bands surrounding the waveforms represent the standard error of the mean (SEM) across participants. The lower panels show the corresponding post-minus-pre stimulation difference waves (Δ) for each electrode and stimulation condition. Dashed vertical lines indicate stimulus onset (0 ms). Cluster-based permutation testing revealed significant differences in pre-to-post ERP changes between stimulation conditions within the posterior region of interest (234–584 ms, p = 0.0002). This significant time window is highlighted by the gray shaded areas in the trough-phase condition, which showed the largest increase in ERP amplitude following stimulation.

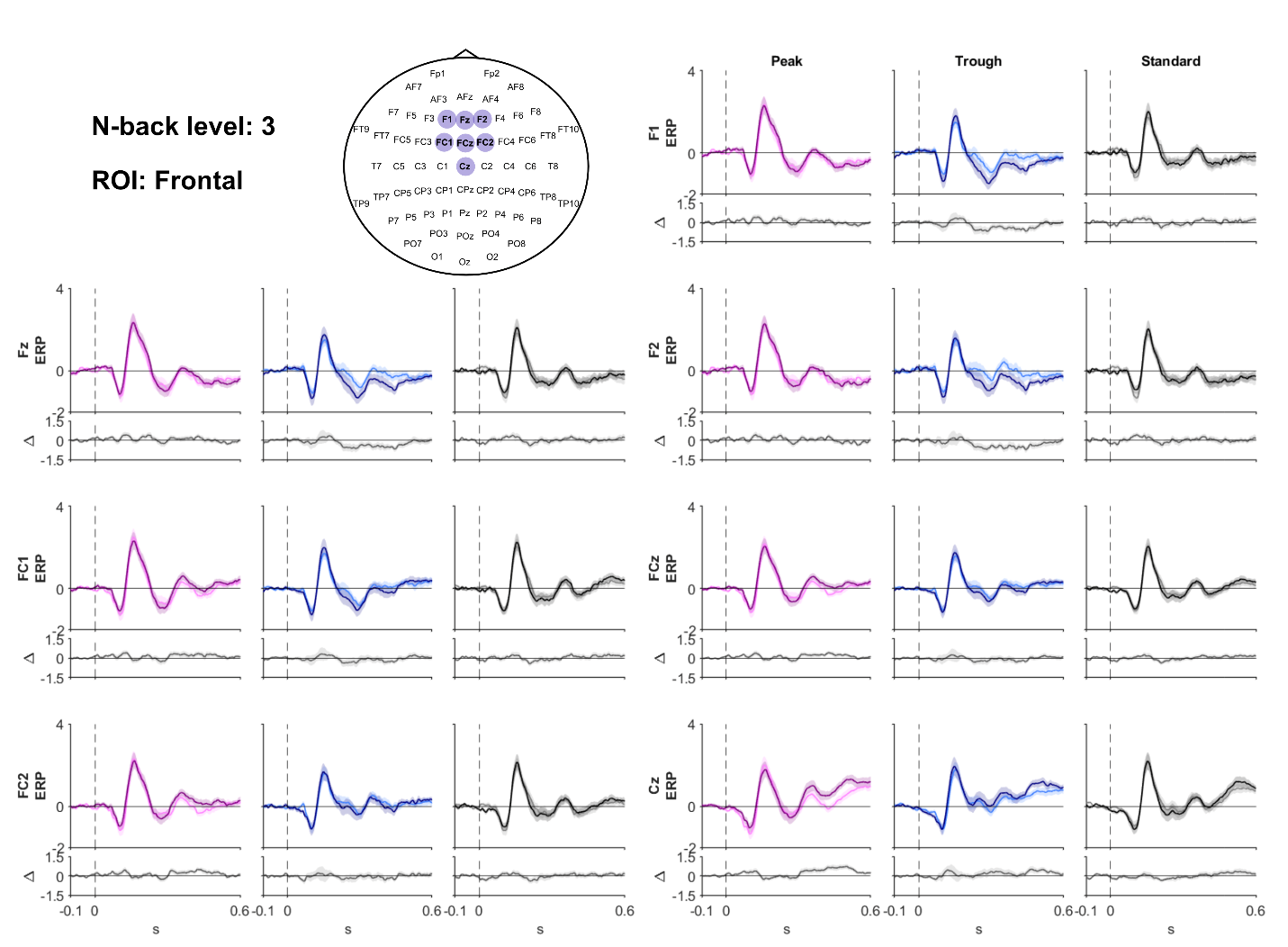

**Supplementary Figure 9. Event-related potentials across frontal electrodes during the 3-back task.** Grand-average ERP waveforms recorded before and after rTMS are shown for peak-phase (magenta), trough-phase (blue), and standard 20 Hz rTMS (black) across seven frontal and frontocentral electrodes (F1, Fz, F2, FC1, FCz, FC2, and Cz), highlighted in the electrode layout. The shaded bands surrounding the waveforms represent the standard error of the mean (SEM) across participants. The lower panels show the corresponding post-minus-pre stimulation difference waves (Δ) for each electrode and stimulation condition. Dashed vertical lines indicate stimulus onset (0 ms). Cluster-based permutation testing revealed no significant differences in pre-to-post ERP changes between stimulation conditions within the frontal region of interest.

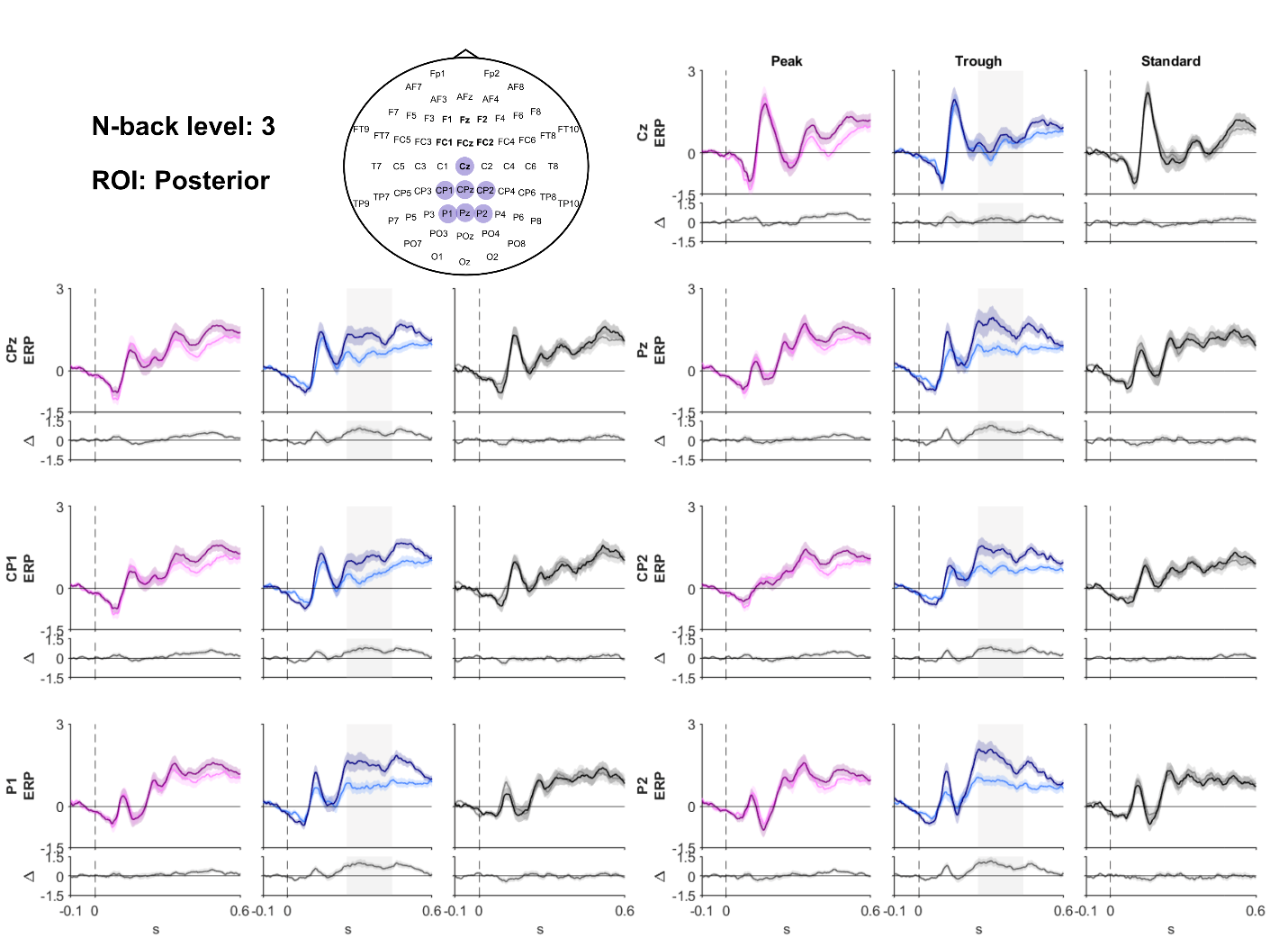

**Supplementary Figure 10. Event-related potentials across posterior electrodes during the 3-back task.** Grand-average ERP waveforms recorded before and after rTMS are shown for peak-phase (magenta), trough-phase (blue), and standard 20 Hz rTMS (black) across seven central and posterior electrodes (Cz, CPz, Pz, CP1, CP2, P1, and P2), highlighted in the electrode layout. The shaded bands surrounding the waveforms represent the standard error of the mean (SEM) across participants. The lower panels show the corresponding post-minus-pre stimulation difference waves (Δ) for each electrode and stimulation condition. Dashed vertical lines indicate stimulus onset (0 ms). Cluster-based permutation testing revealed significant differences in pre-to-post ERP changes between stimulation conditions within the posterior region of interest (246–434 ms, p = 0.0082). This significant time window is highlighted by the gray shaded areas in the trough-phase condition, which showed the largest increase in ERP amplitude following stimulation.

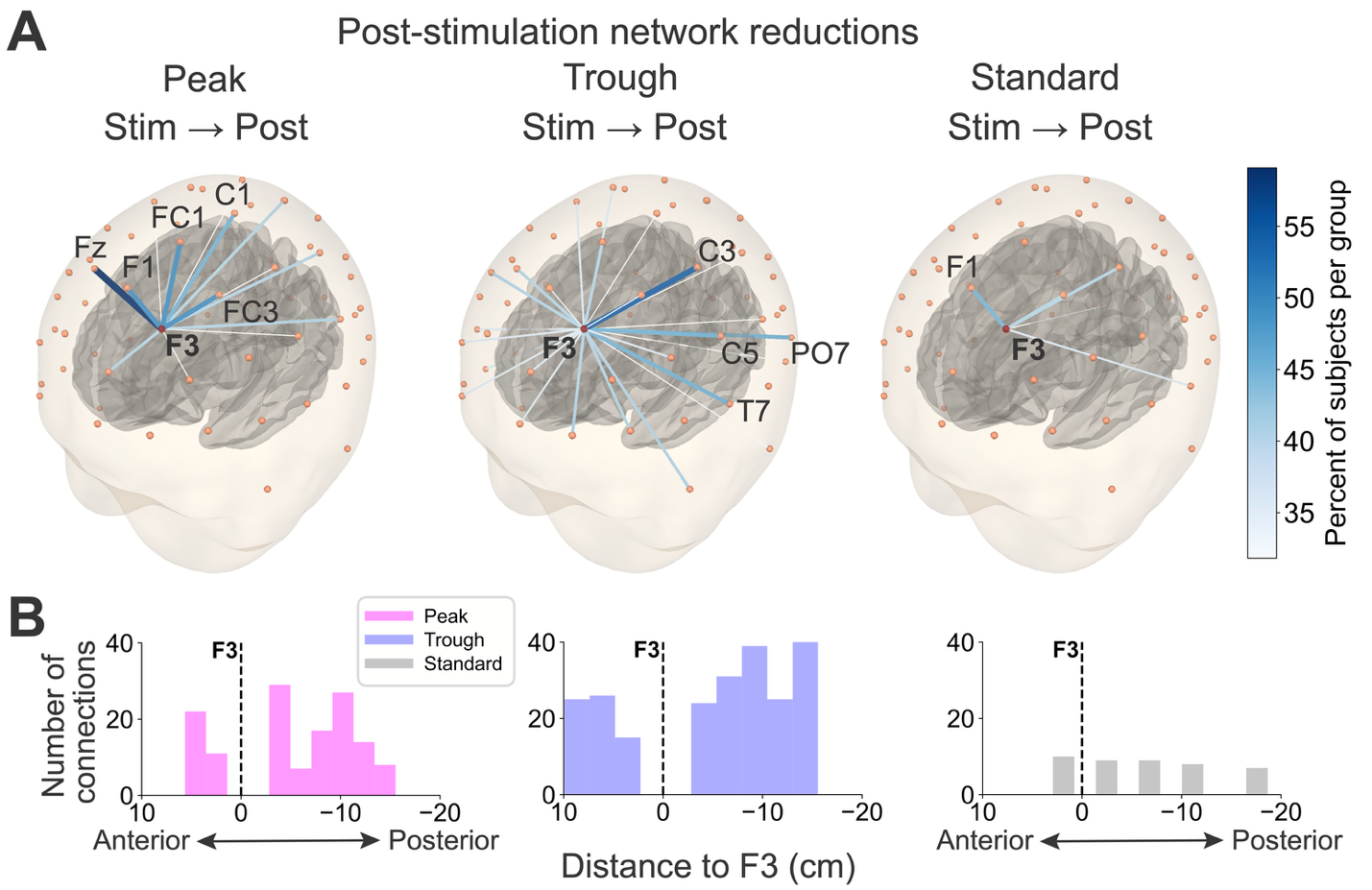

**Supplementary Figure 11. Network size reductions following rTMS.** A) Connections that disappeared from the network centered around F3 from the stimulation block to the post-stimulation block separated by group. All three head models are based on the fsaverage template and show the electrode locations (orange-red spheres) from the montage used in this study on top of the scalp and the gray matter surface below. The color and line thickness of each line represent how many subjects lost this connection from the stimulation to post-stimulation. Labels mark the name and location of F3 and the most commonly removed channel connection for a given group. The connections removed were largely consistent with the ones added from the pre-stimulation block to the stimulation block in Figure 5C. B) Histograms of connections shown in A over signed anteroposterior distances to F3. Colors denote the group. Positive values indicate connections anterior of F3, negative values indicate connections posterior to F3.

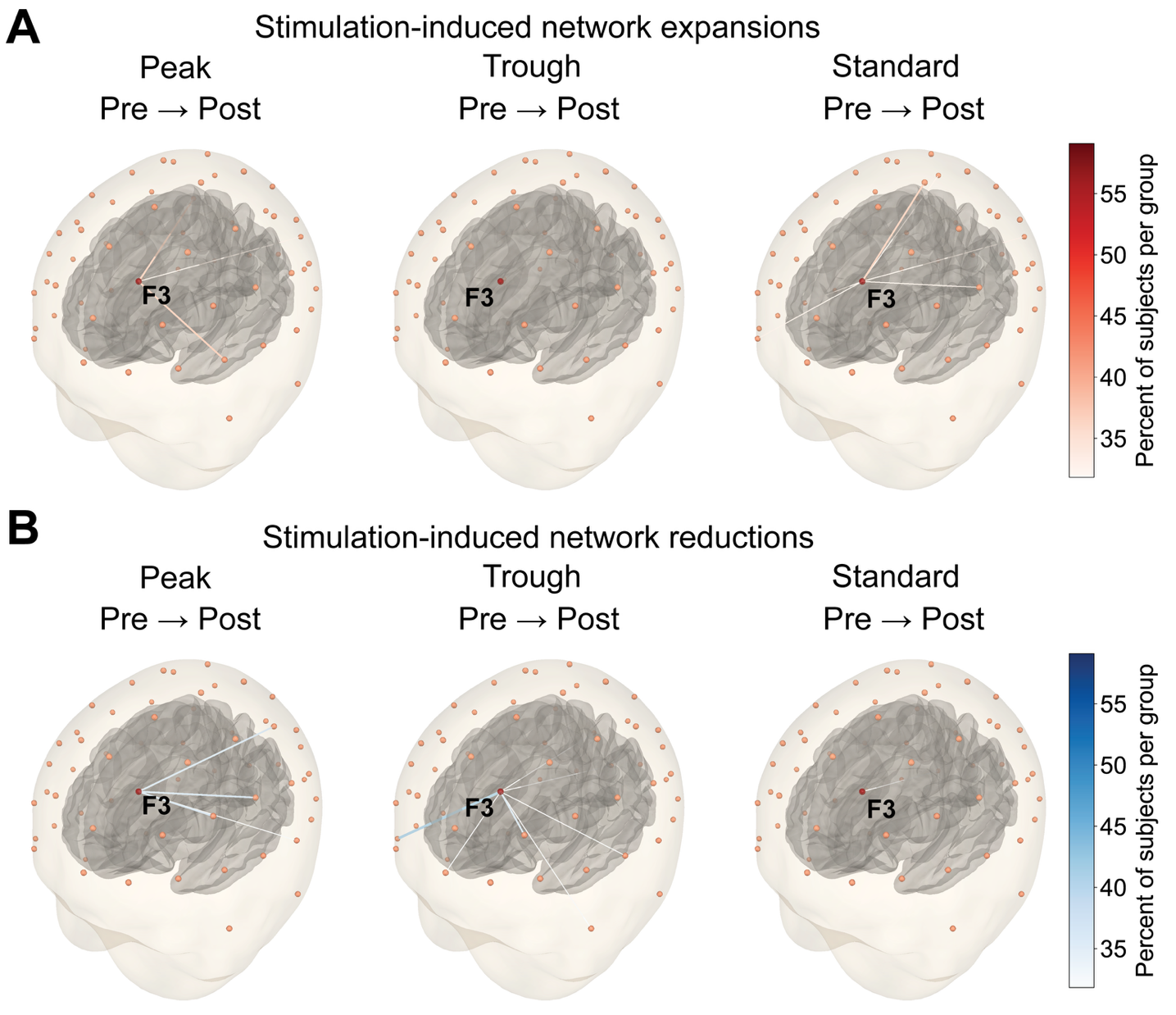

**Supplementary Figure 12. rTMS-induced network changes from pre- to post-stimulation.** A) New connections added to the network centered around F3 from the pre- to the post-stimulation block separated by group. Head models, electrode locations, as well as the line- and color-coding are the same as in Supplementary Figure 11. B) Connections that were removed from the network centered around F3 from the pre- to the post-stimulation block separated by group.
